# Tumor expressed CD95 causes suppression of anti-tumor activity of NK cells in a model of triple negative breast cancer

**DOI:** 10.1101/2021.02.12.431011

**Authors:** Abdul S. Qadir, Jean Philippe Guégan, Christophe Ginestier, Assia Chaibi, Alban Bessede, Emmanuelle Charafe-Jauffret, Manon Macario, Vincent Lavoué, Thibault de la Motte Rouge, Calvin Law, Jacob Vilker, Hongbin Wang, Emily Stroup, Matthew J. Schipma, Bryan Bridgeman, Andrea E. Murmann, Zhe Ji, Patrick Legembre, Marcus E. Peter

**Affiliations:** Division Hematology/Oncology, Department of Biochemistry and Molecular Genetics, Feinberg School of Medicine, Northwestern University, Chicago, IL 60611, USA; Immusmol SAS, 229 Cours de l’Argonne 33000 Bordeaux, France; Aix-Marseille Univ, Inserm, CNRS, Institut Paoli-Calmettes, CRCM, Epithelial Stem Cells and Cancer Lab, Marseille, France; Department of Gynecology, University Hospital of Rennes, Rennes, France; Centre Eugène Marquis, Rennes, France; Department of Medicine, Department of Biochemistry and Molecular Genetics, Feinberg School of Medicine, Northwestern University, Chicago, IL 60611, USA; Department of Pharmacology, Department of Biochemistry and Molecular Genetics, Feinberg School of Medicine, Northwestern University, Chicago, IL 60611, USA; Department of Biomedical Engineering, McCormick School of Engineering, Northwestern University, Evanston, Illinois; INSERM U1262, CRIBL, Université Limoges, Limoges, France

**Author notes:** Shared first authorship (Abdul S. Qadir, Jean Philippe Guégan). Shared last authorship (Patrick Legembre, Marcus E. Peter). Department of Pharmacology, The University of Illinois College of Medicine, Chicago, Illinois, 60612, USA. Contact: Marcus Peter, Patrick Legembre.

**Keywords:** Fas, cancer stem cells, breast cancer, MDSC

## Abstract

The apoptosis inducing receptor CD95/Fas has multiple tumorigenic activities. Stimulation by its cognate ligand CD95L on many cancer cells increases their growth, motility, ability to invade and/or their cancer stemness. Using genetically engineered mouse models of ovarian and liver cancer, we previously reported that deletion of CD95 in the tumor cells strongly reduced their ability to grow *in vivo* [1, 2]. Using a combination of immune-deficient and immune-competent mouse models, we now establish that loss of CD95 in metastatic triple negative breast cancer cells prevents tumor growth by modulating the immune landscape. CD95 deficient but not wild-type tumors barely grow in an immune-competent environment and show an increase in immune infiltrates into the tumor. This growth reduction is caused by NK cells and does not involve CD8^+^ T cells. On the other hand, in immune compromised mice CD95 k.o. cells are not growth inhibited, but they fail to form metastases. In summary, we demonstrate that in addition to its tumor and metastasis promoting activities, CD95 expression by tumor cells can exert immune suppressive activities providing a new target for immune therapy.

## Introduction

CD95/Fas is a well characterized death receptor that in permissive cells or when crosslinked in cancer cells *in vitro* mediates induction of apoptosis when stimulated by its cognate ligand CD95L [3–5]. It is now well established that CD95 also has multiple nonapoptotic functions [6–8]. Many of these activities are tumor promoting (reviewed in [9, 10]). We previously demonstrated that neither a low grade serous nor an endometrial ovarian cancer, nor a chemically (DEN) induced liver cancer efficiently grew in mice after tissue specific deletion of CD95 in the ovaries or the liver, respectively [1, 2]. In fact, in all three models either no tumors formed or their numbers and sizes were severely reduced. Interestingly, all tested DEN-induced liver cancer nodules or low grade ovarian cancer that formed in mice still expressed CD95 [2] indicating that these tumors were escapers caused by the inefficiency of Cre recombination used to delete CD95 and pointing at a pivotal role of CD95 in cancer cell survival *in vivo*. However, it was not clear whether this activity of CD95 was cell autonomous or required cells of the tumor microenvironment.

Among women, breast cancer (BC) is the most common cause of cancer, and the second leading cause of cancer death [11]. BC is a heterogeneous disease whose molecular classification has been significantly improved to distinguish luminal A and B expressing hormonal receptors including estrogen (ER) and/or progesterone receptors (PR), basal/triple negative breast cancer (TNBC), and human epidermal growth factor receptor 2 (HER2)-like tumors. This molecular taxonomy is clinically relevant with basal/TNBC patients presenting the poorest clinical outcome with no targeted therapies available as compared to other molecular subtypes. TNBCs progress rapidly and generate metastases that remain the major cause of cancer-related mortality in these patients [12]. TNBC presents a major intratumoral heterogeneity that contributes to therapy failure and disease progression. The origin of this cellular heterogeneity is mainly explained by several subpopulations of self-renewing breast cancer stem cells (CSCs) sustaining the long-term oligoclonal maintenance of the neoplasm [13]. We recently reported that engaging CD95 on ER positive BC cells contributes to CSC survival [14, 15]. Although CD95L-expressing immune cells edit tumor cells by sparing cancer cells expressing low CD95 level at their plasma membrane [16], CD95 is highly expressed on TNBC [17] and the function of this receptor in these cancers remained unknown.

Myeloid derived suppressor cells (MDSCs) were initially identified in human and mouse cancers due to their potent immunosuppressive activity [18, 19]. MDSCs are a heterogeneous population of immature myeloid cells that include monocytic (M-MDSC, CD45^+^CD11b^+^Gr1^High^Ly6C^High^) and granulocytic (G-MDSC, CD45^+^CD11b^+^Gr1^High^Ly6G^+^) subsets both of which have been shown to exert immune-suppressive activities. Within tumors, additional myeloid cells referred to as tumor-associated macrophages (TAM) display immunosuppressive activity [20].

The presence of these cells inside the tumor seems to favor not only tumor cell survival but also angiogenesis and metastases [21] suggesting that inhibitors of MDSC recruitment inside tumors might represent an attractive therapeutic option [22]. To examine the role of CD95 in a TNBC model, we generated two independent sets of CD95 knockout 4T1 clones using different methods and on different continents. Only in immune competent mice, the deletion of CD95 caused a dramatic loss of tumor growth. This was accompanied by an increased T-cell infiltration (CD4^+^ and CD8^+^ T cells) and a reduction in G-MDSCs inside the tumors. A substantial increase in many other immune cells including natural killer (NK) cells was detected in the k.o. tumors as compared to control cancer cells. Most notably, depletion of tumor-infiltrating macrophages (TAM), CD8^+^ and CD4^+^ T-cells did not restore tumor growth of the k.o. cells while elimination of NK cells did. Our data point at a novel role of CD95 as a general immune suppressive receptor that advanced cancer cells maintain to avoid destruction by multiple immune cells and more specifically, 4T1 tumor cells keep CD95 expression to prevent a NK-driven anti-tumor response.

## Results

### Knock-out of CD95 does not affect the growth or stemness of 4T1 cells *in vitro*

We previously demonstrated that different cancers *in vivo* can barely grow without CD95 expression [1, 2]. In contrast, knocking out CD95 in either ovarian cancer or a BC cell lines did not substantially reduce growth of cancer cells *in vitro* [23]. We were therefore wondering whether the tumor microenvironment and/or a functional immune system was required for CD95 expressing tumor cells to grow *in vivo* or whether they were responsible for the destruction of CD95 deficient tumor cells. Based on the results obtained with our genetically engineered mouse models (GEMMs) in which either tumors did not form or tumors that formed still expressed CD95 due to Cre recombination inefficiency, we concluded that this question could not conclusively be addressed in a CD95 k.o. GEMM. We therefore used CRISPR/Cas9 gene editing to delete CD95 in the aggressive murine TNBC cell line 4T1 that could be grown in syngeneic Balb/c mice [24]. We generated single cell clones with a complete biallelic knockout of CD95. To increase the rigor of the study and to eliminate any possibility of clonal effects or effects of Cas9 expression two sets of k.o. clones were generated starting with different 4T1 lines and using different strategies to generate CD95 deficiency. The U-clones (generated in the US, all experiments in light blue boxes) involved the use of stable expression of GFP-Cas9 and a two guide (sg)RNA system to delete exon 9 of murine CD95 (**Fig. S1A**). On the other hand, the F-clones (generated in France, all experiments in light yellow boxes) were generated by transfecting both Cas9 and a sgRNA-plasmid resulting in a frame shift mutation around the CD95 transcriptional start site (**Fig. S1B**). Two U-clones were confirmed to not contain CD95-exon 9 anymore (**Fig. S1C**). They had reduced total CD95 mRNA expression (**Fig. S1D**), expressed little or no detectable CD95 on their cell surface (**Fig. S1E**), and neither of the two k.o. clones had a consistent up or downregulation of CD95L mRNA when compared to either parental cells or a Cas9 control clone (**Fig. S1F**). Similarly, two k.o. F-clones were selected and expressed no detectable surface CD95 (**Fig. S1G**). Similar to the human cell lines in which we had deleted CD95, none of the four 4T1 CD95 k.o. clones showed growth reduction *in vitro* or major cell cycle changes when compared to their wt counterparts (**Fig. S1H-J**).

We had recently reported for a number of ER^+^ BC cell lines that chronic stimulation through CD95 induced cancer stemness through induction of a type I interferon response which resulted in activation of STAT1 [14, 15]. Stemness could also be increased by treating cells directly with type I interferons. In contrast, in the TNBC cell line 4T1 prolonged stimulation through CD95 or addition of IFNß did not cause a substantial upregulation of CSC driving transcription factors, STAT1 or its major target gene PLSCR1 (**Fig. S2A**). While the stemness marker BMI1 was induced, the two E-box binding proteins ZEB1 and ZEB2 were not. Treatment of 4T1 cells with either LzCD95L or IFNß did also not substantially affect cell growth (**Fig. S2B**) and it did not increase the ability of the cells to form spheres (**Fig. S2C**). It also did not result in an increase in ALDH1 activity (**Fig. S2D**) or upregulation of CD44 (**Fig. S2E**). Finally, treatment of 4T1 cells with LzCD95L did not induce a substantial amount of apoptosis (**Fig. S2F**). These data suggest that cell signaling through CD95 in 4T1 cells is severely impaired despite the fact that these cells express substantial amounts of surface CD95. Although we did not see a significant and reproducible change in the expression of some of the stemness markers in the CD95 k.o. U-clones (**Fig. S3A**), both CD95 k.o. U- and F-clones displayed a reduced ability to form spheres when compared to control cells (**Fig. S3B**, and **C**). In summary, the stimulation of CD95 in 4T1 TNBC cells did not result in a major change in *in vitro* growth or cancer stemness suggesting that CD95 on these cells has low signaling competence. On the other hand, although the stemness markers were not dramatically affected by CD95 loss in 4T1 cells, the loss of this receptor was associated with a slight but significant reduction in the ability of the cells to form spheres, suggesting that the presence of CD95 could alter some tumor features.

### CD95-deficient 4T1 cells grow faster in immune deficient mice than wt cells but do not form lung metastases

To determine whether the lack of CD95 expression would affect growth of 4T1 cells *in vivo,* we carried out an orthotopic graft of luciferase-expressing Cas9 control, or a mixture of the two CD95 k.o. U-clones into the mammary fat pad of NSG mice (**Fig. 1A**). After a lag phase of about 14 days the k.o. U-cells grew more rapidly than wt cells in the NSG mice. When the two k.o. clones and the two wt controls were injected individually tumor weight and volume was similar at two weeks in the mice (**Fig. 1B, C**). The two F-k.o. clones generally grew more rapidly than wt cells (**Fig. 1D**), suggesting that in immune deficient mice, cells without CD95 had a growth advantage. This was not due to a difference in the expression the stemness markers ALDH1 or CD44 (**Fig. 1E**). NSG mice are devoid of lymphocytes and NK cells, but do contain macrophages [25]. While the number of intratumor G-MDSCs was the same in wt and k.o. tumors (**Fig. 1F**), we observed a different distribution of F4/80 or CD163-expressing macrophages in CD95 k.o. as compared to wt tumors (**Fig. S4**). Whereas F4/80 or CD163-expressing macrophages infiltrated wt tumors, these cells remained mostly at the tumor periphery in CD95 k.o. tumors, pointing at a modulation of the immune landscape by tumor expressed CD95 even in NSG mice.

**Figure 1.**
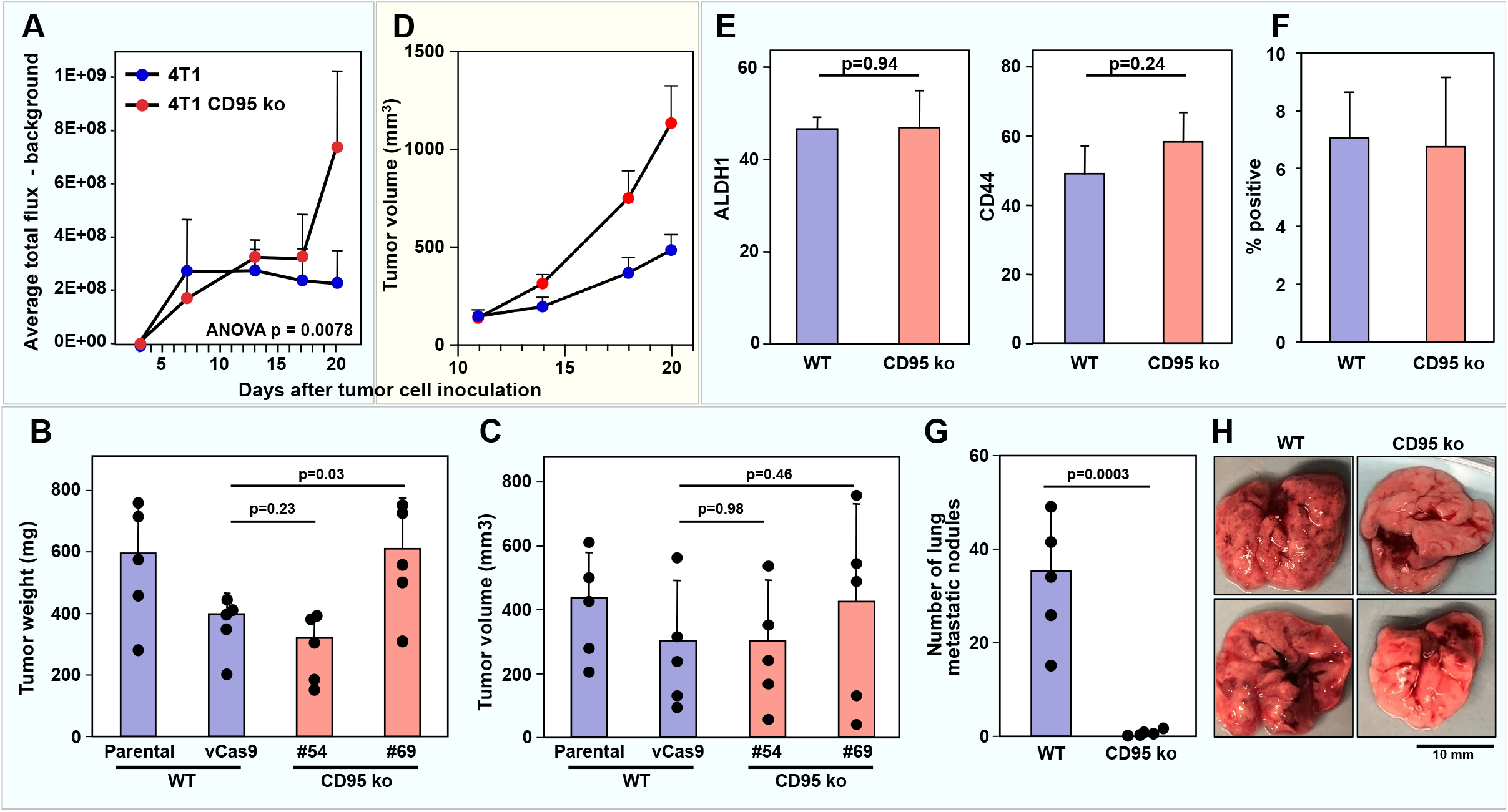
Deletion of CD95 in 4T1 cells does not inhibit tumor growth in NSG mice but blocks lung metastasis. **A** Wild-type (vCas9) or CD95 k.o. 4T1 (mixture of clones #54 and #69) cells (10^5^) were injected into the fat pad of NSG mice. Bioluminescence of tumors was quantified at the indicated times after injection of cells. p-value was calculated using two-way ANOVA. **B, C** Equal numbers of WT (Parental or vCas9) or CD95 k.o. (#54 or #69) 4T1 cells were transplanted into the mammary fat pad of NSG mice, and tumor weight (**B**) and tumor volume (**C**) was measured after two weeks. Data are presented as means ± SEM; n = 5 tumors for each group. **D** Wild-type (mixture of clones #3 and #4) or CD95 k.o. 4T1 (mixture of clones #10 and #12) cells (10^5^) were injected into the mammary fat pad of NSG mice and tumor volume was measured using a caliper. p-value was calculated using ANOVA. **E** ALDH1 activity and CD44 surface staining of single cells suspensions of tumors isolated from NSG mice. **F** IHC quantification of Ly6G positive cells in the wt and CD95 k.o. tumors monitored in B. **G** Equal numbers of WT (Parental or vCas9) or CD95 k.o. (1:1 mixture of #54 or #69) 4T1 cells were transplanted into the mammary fat pad of NSG mice, and tumor nodules of the primary tumor on the surface of the lungs were counted two weeks after transplantation. Data are presented as means ± SEM; n = 5 tumors in each group. **H** The appearance of representative lungs analyzed in G. Experiments involving U-clones are in a light blue box and experiments involving F-clones are in a light yellow box, respectively.

4T1 cells are aggressively growing cancer cells that form lung metastases in implanted mice [26]. While injection of wt 4T1 cells into NSG mice caused multiple lung metastases, barely any lung metastases were detected in mice with implanted CD95 k.o. cells (**Fig. 1G, H**). Interestingly, this occurred despite the fact that the primary CD95 k.o. tumors grew larger than the wt tumors in these mice. Reduced numbers of lung metastases were also seen in NSG mice injected with the F-clones, however, the difference between wt and CD95 k.o. cells was not as dramatic (data not shown). These data are consistent with the known activity of CD95 to increase motility and invasiveness of breast cancer cells [27, 28] and could also be associated with the somewhat reduced capacity of CD95-deficient 4T1 cells to form spheroids as compared to wt counterparts (**Fig. S3B** and **C**).

### CD95 deficient 4T1 cells barely grow in immune competent mice compared to wt cells and show reduced markers of stemness

We next wondered whether the loss of CD95 by tumor cells could impair tumor growth by modulating immune infiltrates. To evaluate this possibility, we orthotopically grafted 4T1 cells into syngeneic Balb/c mice. CD95 k.o. 4T1 cells grew much less than wt cells which only stopped growing when reaching a certain size and becoming necrotic (**Fig. 2A**). The Cas9 control, or a mixture of the two CD95 k.o. U-clones expressed luciferase (**Fig. 2A**). To exclude an immune effect linked to the fact that CD95 wt and k.o. clones expressed the xeno-antigen luciferase [29], we performed the experiment with unlabeled parental cells and compared the growth of parental cells with that of the vCas9 and the two CD95 k.o. U-clones individually (**Fig. 2B, C**). Tumors were harvested after two weeks of growth in the mice. Consistent with the above results, both the parental and unmodified 4T1 cells as well as the Cas9 expressing wt cells grew much faster than the two CD95 k.o. clones as monitored by both tumor volume and tumor weight (**Fig. 2B, C**). In fact, the k.o. cells barely grew in these mice. This behavior of CD95 k.o. cells was largely confirmed by the orthotopic xenograft of the F-clones into Balb/c mice (**Fig. 2D**). Tumor growth of the two k.o. clones was less efficient than that of the parental cells or of a wt clone (**Fig. 2D**). In contrast to the tumors grown in NSG mice, *ex vivo* isolated CD95 k.o. U-clone tumor cells from the Balb/c mice showed a clear reduction of the stemness markers ALDH1 and CD44 when compared to wt tumors (**Fig. 2E**). Unexpectedly, these data suggest that the loss of the death receptor CD95 reduces tumor growth in mice that have a functional immune system.

**Figure 2.**
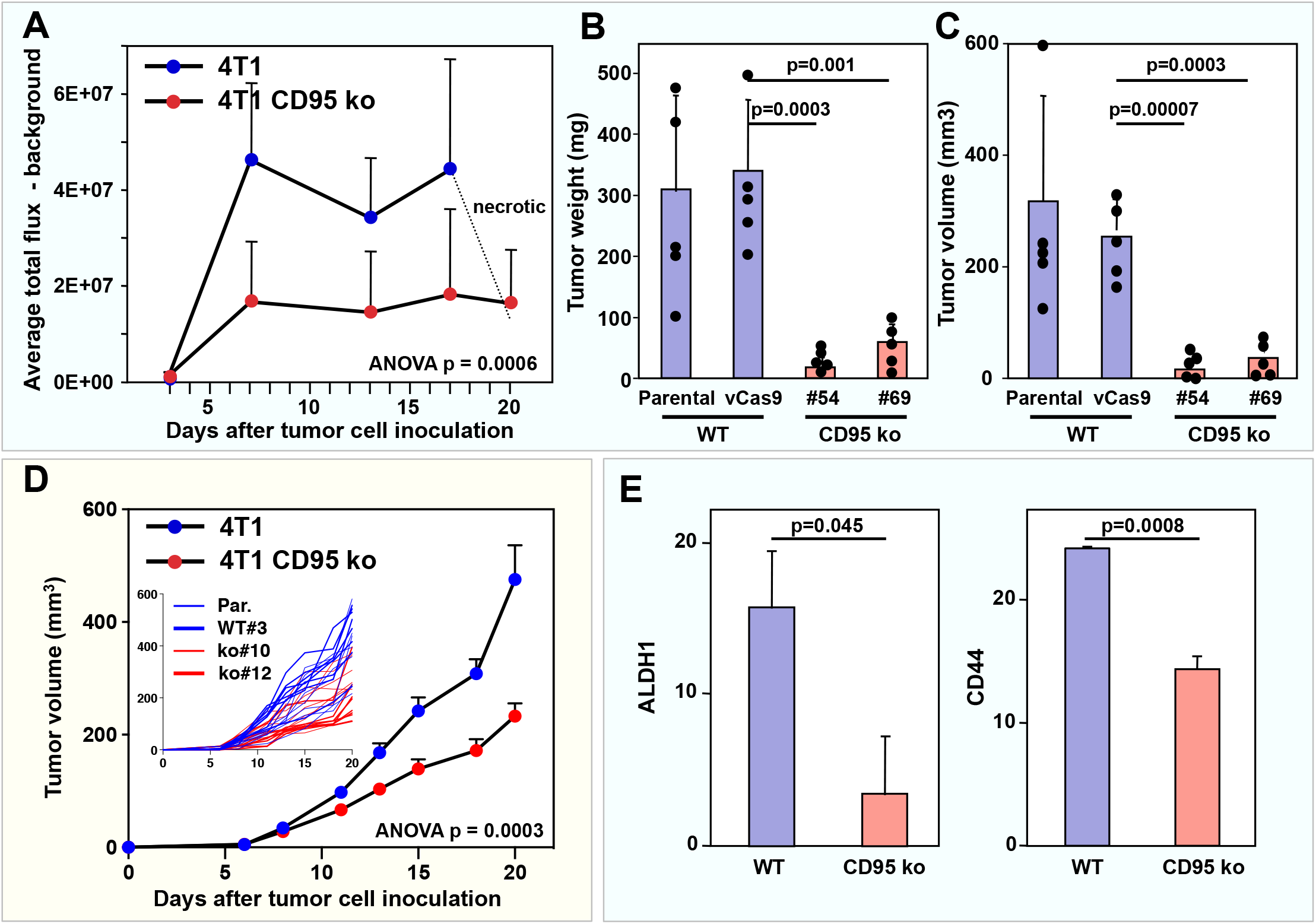
Deletion of CD95 in 4T1 cells inhibits tumor growth and cancer stemness in Balb/c mice. **A** Luciferase expressing wild-type (cCas9) or CD95 k.o. 4T1 (mixture of two k.o. clones) cells (10^5^) were injected into the fat pad of Balb/c mice (5 mice per group). Bioluminescence of tumors was quantified at the indicated times after injection of cells. p-value was calculated using ANOVA. **B, C** Equal numbers of unmodified WT (Parental or vCas9) or CD95 k.o. (#54 or #69) 4T1 cells were transplanted into the mammary fat pad of Balb/c mice, and tumor weight (**B**) and tumor volume (**C)** was measured after two weeks. Data are presented as means ± SEM; n = 5 tumors for each group. **D** Parental, wt (#3) and two CD95 k.o. (#10 and #12) 4T1 cells (10^5^ cells) were injected into the mammary fat pad of Balb/c mice (8 mice per group) and tumor volume was measured using a caliper. p-value was calculated using two-way ANOVA. Insert shows the growth of the individual tumors. **E** ALDH1 activity and CD44 surface staining of single cell suspensions of tumors isolated from BALB/c mice.

### A differential analysis of gene expression changes identifies a general immune activation inside the CD95 k.o. tumors in immune competent mice

To get insights into what is causing the suppression of tumor growth after deletion of CD95, we subjected two 4T1 wt controls (wt and vCas9 expressing) and the two CD95 k.o. U-clones to an RNAseq analysis under three different conditions: cells grown *in vitro*, cells grown in NSG and cells grown in Balb/c mice. As the goal was to identify altered genes in the tumors between wt and k.o. tumor specifically in the Balb/c mice, we performed a differential gene expression analysis using DESeq2 (**Table S1**). The number of significantly expressed genes across all samples was 13,083 and the numbers of genes differentially expressed (padj <0.05) between wt and k.o. cells was 496 (cell lines), 485 (NSG mice), and 620 (Balb/c mice). This amounted to 1090 differentially expressed unique genes. We performed k-means clustering with 6 clusters (determined by elbow plot) and plotted a heat map of adjusted expression to identify trends and unique groups (**Fig. 3A**). The top most significant GO terms are listed for each cluster. Genes in the top two clusters were most highly deregulated specifically in the tumors grown in Balb/c mice when compared to either *in vitro* grown cells or grown in NSG mice. Genes in cluster 1 were upregulated in the k.o. tumors and genes in cluster 2 were downregulated. These changes could either be caused by gene expression changes in the tumor cells or more likely caused by infiltrating immune cells. Most importantly, due to the way the data were analyzed these changes did not reflect the difference between NSG and Balb/c mice but rather the difference between wt and CD95 k.o. tumors in the context of immune competent mice. To identify the function of the most highly deregulated genes, we ranked all deregulated genes across all 6 clusters according to the greatest fold difference in the k.o./wt ratio between Balb/c compared to NSG mice (**Table S2**) and subjected this list to a GO analysis (**Fig. 3B**). 10 of the 22 most significantly enriched GOs in the Balb/c mouse tumors were related to immune function and activation of specific immune cell subsets. This result pointed at a general increase in immune cell infiltration and activity in tumors lacking CD95.

**Figure 3.**
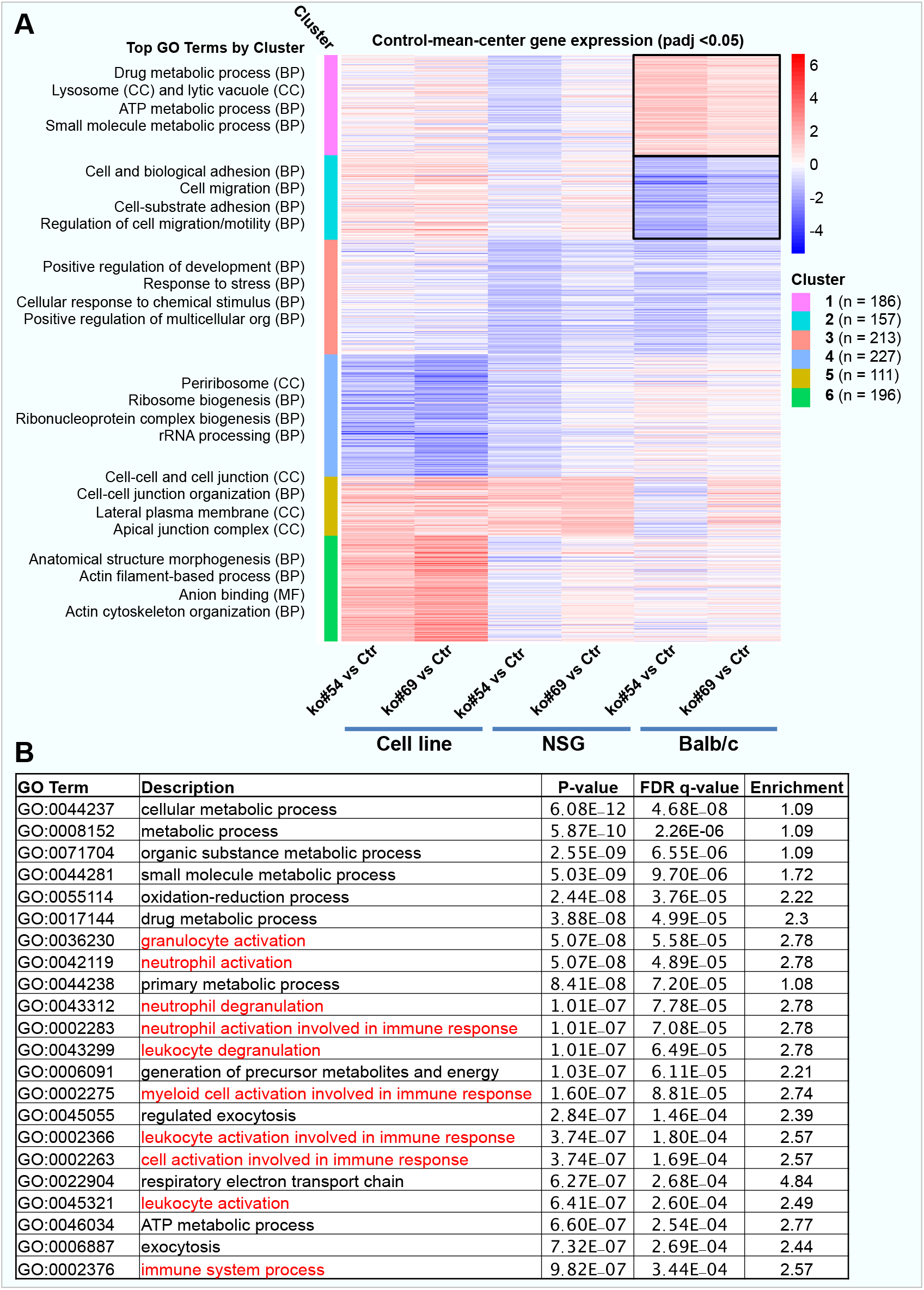
Identification of genes selectively upregulated in the CD95 k.o. tumors grown in immunocompetent mice. **A** Heatmap of control-normalized gene expression for significantly differentially expressed genes adjusted for the average expression in the respective control samples (DESeq2 adjusted *P*-value < 0.05). Genes were clustered using k-means with 6 clusters. The most enriched gene ontology terms for each cluster are shown to the left. **B** GOrilla analysis of a ranked list of the genes differentially expressed in the CD95 k.o. versus the wt cells shown in A. Data were ranked according to the highest difference between the fold upregulation of genes in k.o. versus wt tumors in Balb/c and NSG mice. GO terms associated with immune infiltration and activation are shown in red. Experiments were performed with the U-clones.

### Deletion of CD95 causes an increase in infiltration of most immune cells into the tumor while infiltration of G-MDSCs is reduced

Based on the results of the differential gene expression analysis of CD95 k.o. cells, we subjected U-clones grown as tumors in Balb/c mice to immunohistochemistry analysis (**Fig. 4A**). Macrophages (F4/80 staining) were found mostly at the periphery of the tumors and they were more abundant in CD95 k.o. tumors (**Fig. 4B**). T cells (CD4^+^, CD8^+^ and regulatory T cells [Tregs – FOXP3 staining]) infiltrated k.o. tumors more readily compared to wt tumors. This was pronounced for CD8^+^ T cells which were abundant and showed the most homogenous staining inside the CD95 k.o. tumors, while very few CD8^+^ T cells were detected inside wt tumors (**Fig. 4A, B**). We also detected an increase in phospho-STAT1 in small islets inside the k.o. tumors consistent with an increase in immune infiltration. Tumor vessel density (CD31 staining) was similar between wt and k.o. tumors (**Fig. 4A, B**). These results suggested that CD95 expression in cancer cells could prevent immune cells including cytotoxic killer cells from infiltrating the tumors but had no effect on tumor vascularization, at least not in this rapidly growing tumor model. The IHC analysis of the U-tumors was followed up by a detailed multi-parameter flow cytometry analysis of the tumor infiltrate inside the F-clones grown in Balb/c mice (**Fig. 4C**). This analysis independently confirmed that a large number of different immune cells had preferentially infiltrated the CD95 k.o. tumors including macrophages (CD45^+^CD11b^+^F4/80^+^), both M1 (CD45^+^CD11b^+^F4/80^+^CD38^+^) and M2 (CD45^+^CD11b^+^F4/80^+^CD38^-^) macrophages [30], and all types of T cells as evidenced by an increase in CD3^+^, CD4^+^ and CD8^+^ cells (**Fig. 4C**). Interestingly, while the number of M-MDSCs was increased, the number of G-MDSCs was reduced in the k.o. tumors (**Fig. 4C**). To independently confirm this finding, we stained the U-clone tumors for Ly6G to detect G-MDSCs and confirmed the staining to be strongly reduced in the k.o. tumors (**Fig. 4D** and **4E**, left panel). These findings indicate that in TNBC cells, CD95 expression can prevent tumor infiltration of many cytotoxic immune cells including CD8^+^ T-cells.

**Figure 4.**
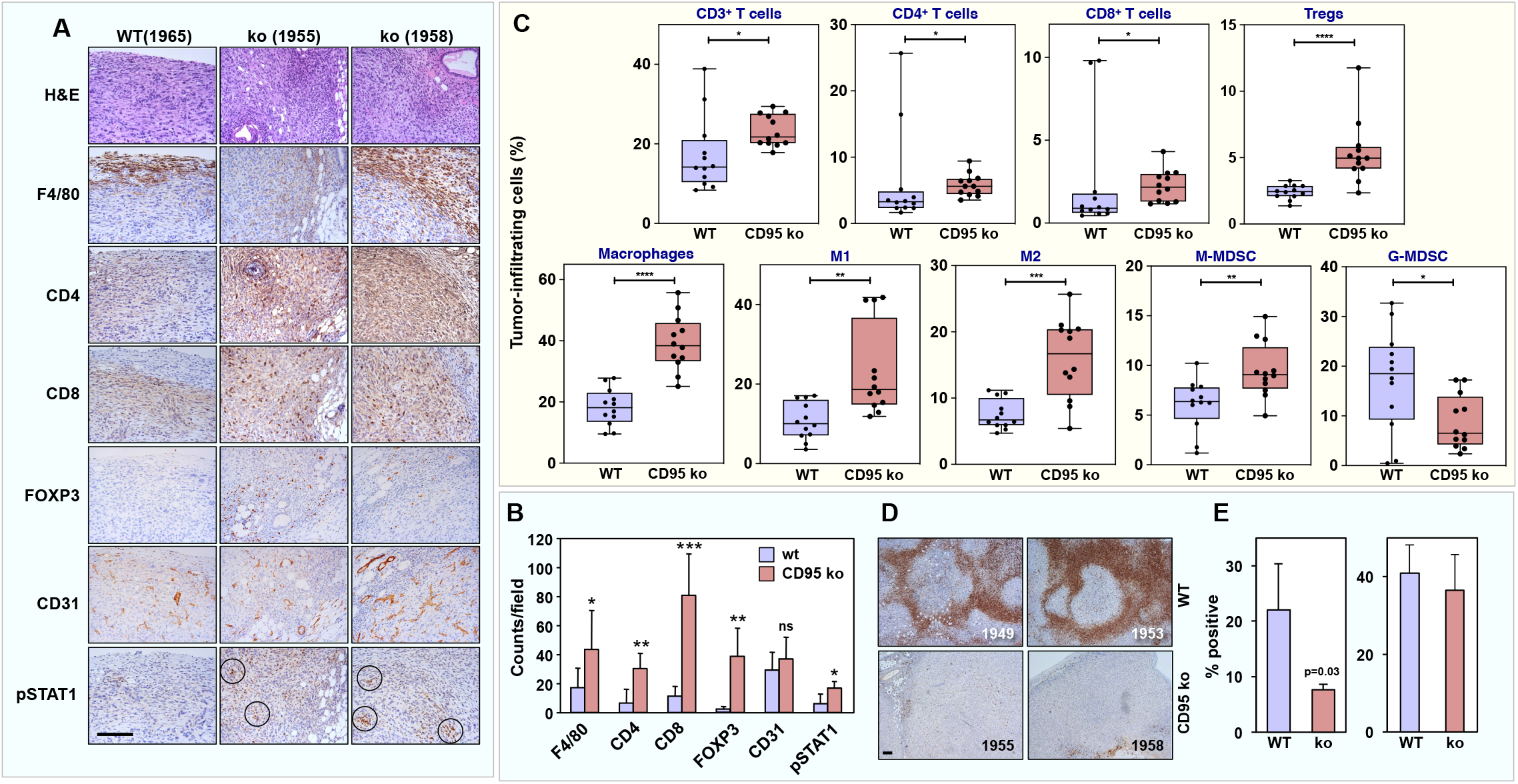
CD95 loss in TNBC cells increases general immune cell infiltration with only G-MDSCs being reduced when compared to wt tumors. **A** WT or CD95 k.o. 4T1 cells were injected into the mammary fat pad of Balb/c mice. Mice were sacrificed when tumors reached maximum allowed size or signs of ulceration were evident. Representative tumor sections of one mouse grafted with wt (1965) and two mice grafted with k.o. cancer cells (1955, 1958) stained with hematoxylin and eosin (H&E) or with antibody specific for mouse macrophages (F4/80), T-cells (CD4, CD8, FOXP3), endothelial cells (CD31), or pSTAT1. Representative images of the immuno-histochemical analysis and H&E staining are shown. **B** The staining intensity of F4/80, CD4, CD8, pSTAT1, CD31and FOXP3 was quantified. Each bar represents the mean ± SEM (*n* = 8). p-value *<0.05, **<0.001; ***<0.0001; ns, not significant. **C** After 21 days, tumors grown in Fig. 2D were resected and dissociated. Tumor-infiltrating cells including macrophages (CD45^+^CD11b^+^F4/80^+^); whole CD3^+^ T-cell population, CD3^+^CD4^+^, CD3^+^CD8^+^ T-cells and Treg (CD3^+^CD4^+^CD25^High^FoxP3^+^), subsets; M1, (CD45^+^CD11b^+^F4/80^+^CD38^+^), M2 (CD45^+^CD11b^+^F4/80^+^CD38^-^); M-MDSC (CD45^+^CD11b^+^Ly6C^+^Ly6G-) and G-MDCS (CD45^+^CD11b^+^Ly6C^Low^Ly6G^+^) were quantified by multiparameter flow cytometry with the indicated combination of markers. **D** Representative tumor sections of two mice injected with wt (1963, 1949) and two mice injected with k.o. cancer cells (1955, 1958) stained for Ly6G. **E** IHC quantification of Ly6G positive cells (left, examples shown in D) and CD11b (right) in the wt and CD95 k.o. tumors shown in B. Scale bars = 50 μm.

### NK but not CD8^+^ or CD4^+^ T cells are responsible for the reduced growth of CD95 k.o. tumors

Multiple studies have shown that tumor infiltrating lymphocytes (TILs) have a strong prognostic impact on women affected by TNBC [31, 32]. A major reduction in disease and distant recurrences was reported for TNBC patients having high amounts of TILs [33]. We first tested to what degree infiltrating CD8^+^ T cells detected in the k.o. tumors were responsible for the reduced tumor growth. To address this question, we depleted CD8^+^ T cells in Balb/c mice by repeated injection of a neutralizing anti-CD8 mAb (**Fig. S5A**). Staining for CD8^+^ T cells in the spleen of depleted mice was undetectable while that of CD4^+^ T cells remained unaffected (**Fig. 5A, B**). 4T1 cells (U-clones) when grown *in vivo* resulted in a substantial increase in spleen size and weight as compared to CD95 k.o. counterparts (**Fig. 5C**). Of note, the reduced growth of CD95 k.o. cells compared to wt cells *in vivo* was not affected by the CD8^+^ T-cell depletion despite the fact that CD8 infiltration was substantially reduced in the CD95 k.o. tumors upon CD8 depletion (**Fig. S6**). CD8 depletion had some effect on tumor growth in general as both wt and k.o. tumors grew slightly larger in the depleted mice (**Fig. 5D**). Consistent with that observation, when areas of the tumors were quantified for TUNEL staining that did not show overt cell death or necrosis (viable tumor tissue), CD8^+^ T-cell-depleted tumors (both CD95 k.o. and wt 4T1 cells) experienced less TUNEL staining than untreated tumors but no difference was observed in fold change between CD95 k.o. and wt tumors (**Fig. 5E**, right panel). These data suggested that although increased CTL infiltration into the k.o. tumors did not explain the strongly reduced tumor growth after deletion of CD95, CD8^+^ T cells did seem to contribute to the increased cell death seen in the 4T1 cells in general.

**Figure 5:**
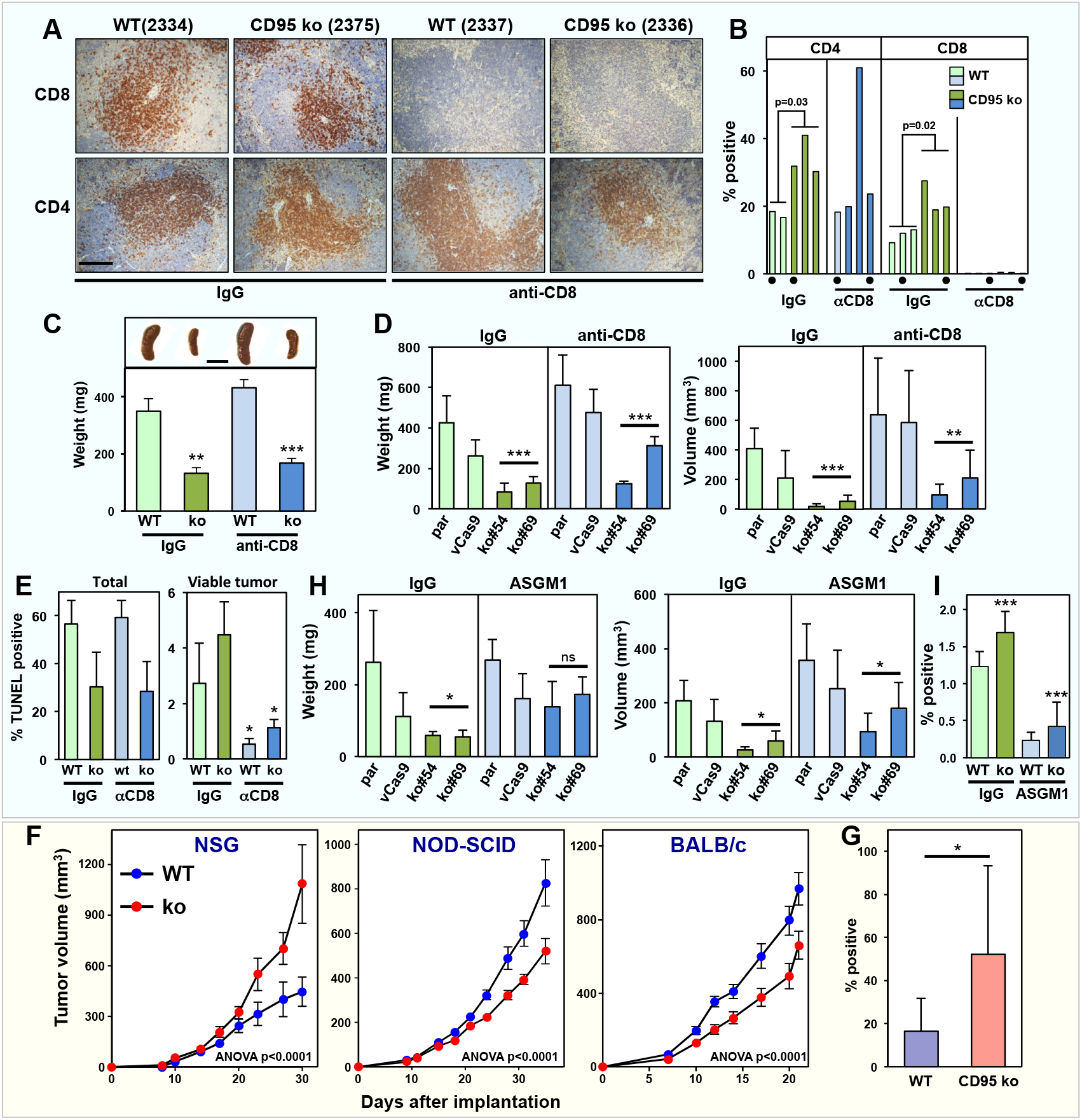
Depletion of NK cells but not of CD8^+^ T cells reverse the growth defect of CD95 k.o. cells in immune competent mice. **A** Representative IHC images of spleens from mice treated with either isotype matched control antibody or anti-CD8 as indicated in A and stained for CD4^+^ or CD8^+^ T cells. Scale bar = 50 μm. **B** Quantification of CD4 and CD8 T cells in the spleen of mice treated with either IgG or anti-CD8. Values labeled by a dot correspond to the images shown in A. **C** *Top*, Representative images of spleens of treated mice. *Bottom*, average spleen weight of treated mice (n=3). Scale bar = 1 cm. **D** Tumor weight (left) and tumor volume (right) in anti-CD8-treated mice (n=5) 16 days after tumor injection. **E** IHC quantification of TUNEL staining of sections of wt and k.o. tumors in mice treated as in D. **F** Tumor growth of a mix of wt 4T1 cells and a CD95 k.o. clone #4 in three different mouse strains. p-values were calculated using two-way ANOVA. **G** IHC quantification of NKp46 positive NK cells in wt and CD95 k.o. tumors grown in NOD-SCID mice shown in F. **H** Tumor weight (left) and tumor volume (right) in anti-Asialo GM1-treated mice (n=5) 16 days after tumor injection. **I** Quantification of NKp46 positive NK cells in the spleen of mice treated with either IgG or anti-Asialo GM1. p-value *<0.05; **<0.001; ***<0.0001. ns, not significant.

In a similar experiment, we confirmed that depletion of CD4^+^ T cells also did not have a significant effect on the reduced growth of CD95 k.o. tumors (**Fig. S5B and S7A, B**). Because of the differences in the amounts of M-MDSCs observed in wt and k.o. tumors (**Fig. 4C**), we treated mice with an antibody to neutralize CSF1-R which has been shown to prevent the recruitment of tumor-infiltrating myeloid cells including macrophages and M-MDSCs [34, 35] (**Fig. S5C**). This experiment was also carried out because among all the CSF ligand and CSF receptor genes in our differential gene expression analysis, CSF1-R was the most highly deregulated gene between the two CD95 k.o. clones and the wt cells, both *in vitro* and when grown as tumors in Balb/c mice (but not in NSG mice) (**Table S3**). Targeting CSF1-R had no effect on tumor growth of either wt or CD95 k.o. tumors despite this antibody treatment leading to the decrease in tumor-associated macrophages (TAMs) and M-MDSCs and an increase in CD4^+^ and CD8^+^ T-cells (**Fig. S7C-E**).

Finally, to test whether NK cells could be responsible for the reduced growth of CD95 k.o. tumors in Balb/c mice, we compared side-by-side the growth of wt and k.o. tumors in NOD-SCID IL2Rgamma^null^ (NSG), NOD-SCID and Balb/c mice. NOD-SCID mice differ from their NSG counterparts mainly in the presence of NK cells [36, 37]. Interestingly, the growth reduction of the CD95 k.o. tumors in NOD-SCID mice was similar to the one seen in Balb/c mice and reversed to the situation in NSG mice (**Fig. 5F**). Consistent with this finding, CD95 k.o. tumors exhibited a significantly higher number of tumor-infiltrating NK cells as compared to wt tumors in NOD-SCID mice (**Fig. 5G**). This was also seen on the mRNA level in a NanoString analysis of wt and k.o. tumors grown in Balb/c mice (data not shown). Finally, depletion of NK cells in Balb/c mice by repeated injections of an anti-Asialo GM1 Ab (**Fig. S5D**), significantly minimized the growth difference between wt and k.o. tumors (**Fig. 5H, I**).

In summary, our data emphasized that in a mouse model of TNBC, the expression of CD95 on tumor cells regulated the global immune landscape. When CD95 was lost in the tumor cells while the total number of immune cells was reduced (as evidenced by the strong reduction in spleen size), the percentage of TILs increased with NK cells being responsible for the anti-tumor response.

## Discussion

We previously reported that deletion of CD95 *in vivo* strongly reduced tumor formation and growth in three different mouse tumor models [1, 2]. In all cases, endogenous CD95 was deleted using Cre recombinase. In two of these models, one involving DEN induced liver cancer and one low grade ovarian cancer, we determined that most if not all tumor nodules that grew in these CD95 k.o. mice still expressed CD95 [1, 2] due to inefficient deletion by Cre. In the current work, we therefore used a syngeneic orthotopic cell line model that allowed us to selectively and completely delete CD95 in all cancer cells and assess its effect on tumor growth. We chose BC as a model because we had previously identified CD95 as a driver of cancer stemness in the context of BC [14, 15, 38]. We found that although the loss of CD95 in TNBC cells did not substantially affect cancer cell survival in an immune-depressed mouse model, when introduced into immune competent mice CD95 k.o. tumors barely grew.

A recent analysis of a cohort of 667 BC patients revealed that CD95 expression increases with advancing grade of disease [17]. In fact, ER-negative BC had almost twice as much CD95 expression as ER-positive BC and TNBC more than twice as much expression. The TNBC 4T1 cell line is part of a series of five BC cell lines that were isolated from mice after sequential growth and differ in their metastatic potential [39]. 4T1 is the most aggressive of these cell lines. Consistent with the data on human TNBC, 4T1 cells have the highest expression of CD95 mRNA of all five isolates (data not shown).

It was previously shown that CD95 signal-mediated inflammation is involved in promoting progression of 4T1 TNBC cells [40] through recruitment of MDSCs which have been reported to drive the growth of 4T1 tumors [41, 42]. We now demonstrate that the complete deletion of CD95 in 4T1 cells results in tumors that barely grow in immune competent mice but have no impairment to grow in immune deficient mice. Our differential analysis of genes upregulated in the k.o. tumors suggested a fundamental recruitment of immune cells with antitumor activity as multiple mRNAs were detected consistent with both leukocyte and neutrophil degranulation (see Fig. 3B). The strongly decreased recruitment of G-MDSCs together with the increase in CD8^+^ T and NK cells in the CD95 k.o. tumors we then observed, left two possibilities to explain the dependence of tumor cells on CD95 to grow *in vivo*: 1) Wt tumors grow more because MDSCs are driving their growth and the increased infiltration of CD8^+^ T cells in k.o. tumors might be the result of the absence of MDSCs. 2) The reduced growth of k.o. tumors might be due to an increased recruitment of TILs. Interestingly, a good prognostic score was shown to be associated with CD8^+^ TILs in ER-negative disease with high CD95 expression [17] suggesting that CTLs attack high CD95-expressing tumor cells. However, we now demonstrate that although CD8^+^ CTLs exert a general anti-tumor effect in the 4T1 model, the loss of CD95 in these TNBC cells is associated with the recruitment of NK cells that reduced tumor growth, ruling out that it is only the MDSCs that drive the growth of wt cells. This finding is consistent with a report showing that treatment of mice with IL-21, a cytokine involved in the NK cell proliferation and activation [43], enhances 4T1 tumor rejection [44]. Not much is known about the role of NK cells in TNBC but molecular signatures associated with NK cells has been shown to be predictive of relapse free survival in BC patients including TNBC patients [45].

In conclusion, we can now separate different observations on the apoptosis independent and cancer relevant functions of CD95 into at least four activities. 1) CD95 drives metastasis formation in a cell autonomous fashion. This observation is consistent with multiple reports demonstrating increased motility or invasiveness of tumor cells upon CD95 stimulation [27, 28, 40, 46–53]. Interestingly, we found that 4T1 cells only form metastases when CD95 is expressed, suggesting that it is indeed a metastatic driver. This activity does not require lymphocytes as this feature was also found in NSG mice. 2) Stimulation of CD95 on cancer cells increases their proliferation. We reported evidence of increased CD95 driven proliferation of cancer cells and in fact stimulating apoptosis resistant cancer cells through CD95 increases their growth rate [54–56]. 3) CD95 stimulation increases cancer stemness [38, 57]. We previously showed that chronic stimulation of CD95 on multiple cancer cell lines including ER^+^ BC induces production of type I interferons, which through activation Type I IFN receptors and activation of STAT1 drives cancer stemness [14, 15]. 4). In the present study, we uncover a novel and unexpected activity of CD95 as an immune suppressive receptor that prevents infiltration of multiple immune cells into the tumor.

We do not believe that the findings on the role of CD95 as a suppressor of tumor immune infiltration is limited to the 4T1 TNBC model. Our data provide an explanation for a long standing enigma of why in GEMMs of endometrioid, low grade ovarian cancer, or liver cancer tumors could not efficiently grow *in vivo* after knock-out of CD95 [1, 2]. In addition, in the model of low grade ovarian cancer, we noticed that the deletion of CD95 in the tumors in the ovaries led to a massive amount of hemorrhage, necrotic cell death and an increase in infiltration of proinflammatory immune cells [2]. We had interpreted this result as a requirement for CD95 for the cancer cells to survive but in this mouse model, we could not determine whether the increase in inflammation we observed was the cause or the consequence of dying tumor cells. Our data now suggest that it was the deletion of CD95 on the developing tumor cells that triggered massive influx of immune cells.

An open question is whether CD95L is required for the immune suppressive activities of CD95. The inability of 4T1 cells to significantly respond to even prolonged CD95 stimulation suggests an activity of CD95 that is independent of its stimulation by CD95L. We can exclude 4T1 cells as a source of CD95L as we did not find a significant number of reads derived from CD95L in our analysis of the parental cells, the wt or CD95 k.o. U clones, consistent with published data [58]. Our data provide support to the idea to not only target CD95L to block the tumor promoting activities of CD95L [59–61] but also to target the CD95 receptor directly.

In summary, this study reveals that CD95 expression on tumor cells impairs the immune attack and when removed enhances the destruction of tumors by the immune system, suggesting that CD95 is an immune checkpoint regulator, and the generation of CD95-targeting antagonists could mimic or add to the therapeutic effects of established checkpoint inhibitors.

## Materials and Methods

### Cell Line and Reagents

The mouse breast cancer cell line to generate the U-clones 4T1 was purchased from the ATCC and cultured in RPMI 1640 medium (Mediatech Inc) containing 10% heat-inactivated fetal bovine serum (FBS) (Sigma-Aldrich), 1% L-glutamine (Mediatech Inc) and 1% penicillin/streptomycin (Mediatech Inc). The 4T1 cell line to generate the F-clones was purchased from ATCC (Molsheim Cedex, France) and cultured in RPMI supplemented with 8% heat-inactivated FCS (v/v) and 2 mM L-glutamine at 37°C/5% CO_2_. IFNβ (11415-1) (used at 1000U/ml) was purchased from pbl Assay Science. Leucine zipper tagged CD95L (LzCD95L) was used to induce apoptosis at 100 ng/ml as described before unless otherwise specified (Qadir et al., 2017). Propidium iodide (#P4864), bovine serum albumin (BSA), EdU flow cytometry kit 488, puromycin, G418 and hydrocortisone were purchased from Sigma-Aldrich. Blasticidin was purchased from InvivoGen. Reagents used for flow cytometry, anti-CD44-APC conjugated (17-0441-82), Isotype Rat IgG2b control (17-4031-82), PE conjugated-anti-CD95 (2-0951-83) and PE conjugated Isotype control (12-4714-82) were from eBioscience. The Aldefluor kit (Stem Cell Technologies) was used for the staining and quantification of ALDH1 activity by flow cytometry. Gates were based on the analysis of cells treated with the specific inhibiting reagent DEAB as a negative control following the instructions by the manufacturer. For the immune histochemistry (IHC) of tumor slides samples the following primary antibodies were used F4/80 (eBioscience™ # 14-4801), CD4 (eBioscience™ #14-9766-82), CD8 (eBioscience™ #14-0195-82), pSTAT1 (Cell Signaling #9167), CD31 (Santa Cruz #sc-1506), Ly6G (BD Bioscience #551459), CD11b (Abcam #ab133357), FOXP3 (eBioscience™ #14-5773-82), and NKp46 (R&D Systems, AF2225). The secondary antibodies Biotin-SP-AffiniPure donkey anti-rabbit IgG (H+L) (#711-065-152) and Biotin-SP-AffiniPure donkey anti-rat IgG (H+L) (#712-065-153) were from Jackson ImmunoResearch Laboratories.

### CRISPR/Cas9 mediated deletion of CD95

Generation of the CD95 k.o. U-clones: In order to generate CD95 k.o. 4T1 clones, first stably expressing lentiCas9-Blast-4T1 cell (vCas9-4T1) lines were established. The lentiCas9-Blast virus was obtained from the viral core facility of Northwestern University. 4T1 cells (10^5^/well) were plated in 6 well plates and on the next day cells were infected with the indicated lentiCas9-Blast virus supernatants with polybrene (final concentration: 8 μg/ml). Culture supernatants were replaced with fresh medium containing antibiotics 48 hours after transduction. Uninfected cells were eliminated by a selection medium RPM1-1640 including 4 μg/ml Blasticidin. The two guide RNAs for exon 9 deletion of CD95 gene GCCGGGTTCACATCTGCGCTAGG and CTTGAGGAATCTGATGACCGTGG were designed using the CRISPR design tool available at crispr.mit.edu. The 19-20nt crisprRNA (crRNA) complementary to the target of interest and 67nt transactivating crRNA (tracrRNA), which served as a bridge between the cas9 protein and the crRNA, were ordered from IDT. vCas9-4T1 cells were transfected with crRNA and tracrRNA according the manufactures protocol using TransIT-X2 transfecting reagent (Mirus) in 12 well plates. 3 days after transfection cells were subjected to cell sorting at 1 cell per well directly into 96-well plates. After two to three weeks, single cell clones were expanded and subjected to genotyping and single cell cloning. Deletions in clones were verified by genomic PCR using external and internal primers pairs. To detect the Exon 9 specific deletion in the CD95 gene, external primers were 5’-GGAACAGACCAAGCTTCCGA-3’ (For. primer) and 5’-TCCAGAACACAGCCATGGTT-3’ (Rev. primer), and two internal reverse primers were 5’-TTGAGTAAATACATCCCGAGAATTG-3’ (For. primer) and 5’-AGCAAGACAACAGAGCAATAGAAAT-3’(Rev. primer) (product size 790bp) and 5’-ATGCATGACAGCATCCAAGA-3’ (For. primer) and 5’-TGCTGGCAAAGAGAACACAC-3’(Rev primer) (product size 338bp). After screening clones, Sanger sequencing was performed to confirm the proper deletion had occurred.

Generation of the CD95 k.o. F-clones: The following sgRNA sequences were cloned within PX459-V2 plasmid and then transfected with lipofectamine 3000 in 4T1 to generate cell lines deficient for CD95 according to manufacturer’s instructions. sgRNA sequences target mouse CD95: 5’CACCGCTGCAGACATGCTGTGGATC3’ (Fwd), 5’AAACGATCCACAGCATGTCTGCAGC3’ (Rev). After transfection and puromycin selection for 48h, genome-edited cells were cloned by limited dilution and CD95-deficient cells were selected by flow cytometry.

### FACS analysis

For CD95 surface-staining cell pellets of about 10^5^ cells were resuspended in 100 ml of PBS on ice. After resuspension, 5 μl of either anti-CD95 PE conjugated primary antibody or matching isotype control were added. Cells were incubated on ice at 4°C, in the dark, for 30 min, washed twice with PBS, and percent CD95 positive cells were determined by Becton Dickinson LSR Fortessa at the Flow Cytometry Core Facility of the Northwestern University. ALDH1 activity was quantified as previously described (Ceppi et al., 2014). For CD44 staining, cells were washed twice with PBS and stained with primary antibody or isotype control for 30 min at 4°C in the dark, washed twice with PBS and CD44-APC positive cells were quantified. For the analysis if *ex vivo* tumor, wt and CD95 k.o. 4T1 tumors were extracted and minced with sterile razor blades and incubated for 2 h in the presence of Collagenase (1,000 units per sample) in sterile Epicult media (Stem Cell Technology). Cells were washed with sterile filtered PBS supplemented with 1% bovine serum albumin (PBS-BSA 1%) and filtered through a 40 μm nylon mesh (BD Biosciences). For the detection of ALDH1 or CD44, cells were stained as describe above. Data were analyzed using FlowJo version 8.8.6 (Treestar Inc).

To analyze immune cells infiltrating F-clone in Balb/c mice, tumors were harvested, dissociated on gentleMACS using the tumor dissociation kit (Miltenyi Biotec) and cells (250,000 cells) were stained with the following antibody panels. T lymphocytes-infiltrating tumors were identified using an antibody panel consisting of CCR4-BV421, CXCR5-BV605, CD8-BV650, CXCR3-BV711, CD45-BV785, CD4-AF488, CD3-PerCP/Cy5.5, CCR6-PE/Dazzle594 antibodies and Zombie NIR for cell viability. MDSC and macrophage-infiltrating tumors were stained using CD11b-BV510, F4/80-BV605, CD45-BV785, Gr1-PE/Dazzle594, Ly6C-PE/Cy7, Ly6G-APC antibodies and Zombie NIR for cell viability. M1/M2 cells and Treg cells were monitored in permeabilized cells using FoxP3-BV421, CD11b-BV510, F4/80-BV605, CD45-BV785, CD4-AF488, CD3-PerCP/Cy5.5, CD38-PE-Dazzle594, CD25-PE/Cy7, and Zombie NIR for cell viability. All antibodies used for tumor-infiltrating immune cells came from Biolegend (San Diego, CA, USA). Data were acquired using a Novocyte cytometer (ACEA Biosciences) and analyzed using Novoexpress software.

### Sphere forming assays

The single cell sphere formation assay was performed as previously described (Qadir et. al 2017), In brief, cells were pre-treated with LzCD95L, IFNβ, or received no treatment for 6 days. Cell suspensions were passed through a 40 μm sterile cell strainer (Fisher Scientific) to obtain single cells. The strained cell suspension was serially diluted in Mammocult media (Stem Cell Technology) and seeded at 1 cell/well in ultra-low adherence round bottom 96-well plates (Corning) in triplicate. The cells were cultured in Mammocult media supplemented with 4 μg/ml Heparin (Stem Cell Technology), 0.5 μg/ml hydrocortisone and 10% Mammocult Proliferation Supplement (Stem Cell Technology). After 6-7 days, cultures of U-clones were scanned using IncuCyte Zoom and spheres were counted. For F-clones, spheroids were counted manually. To assess the differences between parental and CD95 k.o. 4T1 cells, the cells were seeded without any treatment.

### Cell growth/proliferation assay

To monitor cell proliferation in parental, vCas9, CD95 k.o. #54 and CD95 k.o. #69 4T1 cells (U-clones), cells were seeded at 1000 cells/well in 96 well plates. Cell growth was monitored in quadruplicate samples at various time points in an IncuCyte Zoom using a confluency mask. To monitor proliferation of 4T1 parental cells with LzCD95L or IFNβ, 1000 cells/well were plated in 96 well, with media containing LzCD95L or IFNβ or no treatment and cell growth was monitored over time.

For F-clones, EdU (5-ethynyl-2’-deoxyuridine) proliferation assay (Sigma-Aldrich) was carried out according to the manufacturer’s instructions. Briefly, 4T1 cells were incubated for 1h with 10 μM of the thymidine analog EdU, then washed and fixed with PFA (4%) for 15 min. DNA synthesis (S phase) is associated with EdU incorporation. Finally, iFluor-488, is added and a ‘click’ chemistry reaction allows its covalent cross-linking to the EdU. Cell proliferation (EdU positive cells) was quantified using the Novocyte flow cytometer.

### Cell death assays and cell cycle analysis

To quantify cell death, 50,000 cells were plated in 12 well plates in triplicate and then either left untreated or treated with different concentration of LzCD95L for 24 hr. The total cell pellet consisting of live and dead cells was either re-suspended in lysis buffer (0.1% sodium citrate, pH 7.4, 0.05% Triton X-100, 50 μg/ml propidium iodide), and after incubating for 2-4 hours in the dark at 4°C, percent cell death was quantified by flow cytometry or cells were re-suspended in media and an equal volume of Trypan blue solution (Lonza) was added. Both living and dead (blue) cells were counted on a hemocytometer under a light microscope. To perform cell cycle analysis, 600,000 cells were plated in 6 well plates in triplicate. After 16-20 hours (at ~80% confluence), plates were gently rinsed with PBS and trypsinized to obtain cell pellets. Cell pellets were then washed (resuspended in 2.5 ml PBS, centrifuged at 500 x g for 5 minutes, and then decanted) and resuspended in 500 μl lysis buffer (see above) and kept on ice and protected from light. Immediately before FACS analysis, samples were spiked with 1:500 DAPI solution as a counterstain to assess cell viability (with corresponding no DAPI controls). Samples were then analyzed on a BD FacsAria SORP 6-Laser Cell sorter at the RHLCCC Flow Cytometry Core Facility. Subsequent data were analyzed using FlowJo 10 software.

### Real-time PCR

Real-time PCR was performed as described recently (Qadir et. al 2017). In brief, total RNA was extracted using QIAzol Lysis reagent (Qiagen Sciences) and RNA concentration was measured using a NanoDrop 2000. 1-2 μg of total RNA was used to generate cDNA using the High-Capacity cDNA Reverse Transcription Kit (Applied Biosystems). Gene expression in mouse cells was quantified using specific primers from Life technologies for mGAPDH (Mm99999915_g1), Exon9 specific mCD95 (Mm01204974_m1), Exon 1-2 specific mCD95 (Mm00433237_m1), mCD95L (Mm00438864_m1) mSTAT1 (Mm01257286_m1), mPLSCR1 (Mm01228223_g1), mBMI1, (Mm03053308_g1), mZEB1 (Mm00495564_m1) and mZEB2 (Mm00497196_m1) using of the comparative ΔΔCT method and expressed as fold differences.

### Growth of 4T1 cells in NGS and Balb/c mice

U-clones: 4T1 vCas9 (wt) and CD95 k.o. clones #54 and #69 were infected with pFU-L2G luciferase lentivirus as previously described (Ceppi et al., 2014). Then cells (1×10^5^) of wt or a 1:1 mixture of the CD95 k.o. clones were injected in 100 μl PBS and Matrigel (Cat#3432-010-01, Trevigen) (1:1 ratio) into the fourth mammary fat pad at the base of the nipple into female NOD-scid-gamma (NSG) mice and Balb/c mice (Jackson lab) according to the Northwestern University Institutional Animal Care and Use Committee (IACUC)-approved protocol. Growth of tumors was monitored weekly by using an IVIS Spectrum *in vivo* imaging system and luminescence was quantified at the regions of interest (ROI = the same area for each mouse encompassing the entire mammary gland) using the Living Image software. For some experiments L2G luciferase (Lucneo) infected cells were further infected with NucLight Red Lentivirus Reagent (EF-1 alpha promoter, Puromycin selection, Essen Bioscience) and the stable cells were individually injected into the fourth mammary. In some cases unmodified cells were injected. In other experiments 4T1 CD95 k.o. #54 cells (1×10^5^) (Luc-neo/Nuc-red) and CD95 k.o. #69 cells (Luc-neo/Nuc-red) were injected into same mouse. To grow F-clones *in vivo:* 5×10^4^ cells in PBS were injected in the fourth mammary fat pad at the base of the nipple into female Balb/c (Charles River, France). From Day 6, tumor volume was measured with a set of calipers and calculated by using the following formula: volume = (length×width2)/2. A colony of immunocompromised NSG mice (NOD/SCID/IL2rγnull) was maintained in house under aseptic sterile conditions. The study was conducted under EU and French animal welfare regulations for animal use in experimentation (European Directive 2010/63/EU and French decree and orders of February 1st, 2013) and approved by local ethics committee (Agreement no. #16487-2018082108541206 v3). After two weeks of injection, tumor volume was measured with a set of calipers and calculated by using the following formula: volume = (length×width^2^)/2.

### Depletion of immune cells in mice

To deplete CD8 T cells mice (5 mice/group) were injected i.p. with 200 μg of either anti-mouse CD8 mAb (clone 53-6.7, BioXCell) or IgG2a isotype control (2A3, BioXCell) both in 100 μl. To deplete CD4 T cells mice (5 mice/group) were injected i.p. with 200 μg of either anti-mouse CD4 mAb (GK1.5, BioXCell) or IgG2b isotype control (2.43, BioXCell) both in 100 μl. To deplete NK cells mice (5 mice/group) were injected i.p. with 50 μg of either anti-Asialo GM1 rabbit polyclonal antibody (ThermoFisher) or polyclonal rabbit IgG (ThermoFisher) both in 100 μl. To inhibit CSF1-R mice (5 mice/group) were injected i.p. with 20 mg/kg of either anti-mouse CSF1-R mAb (clone AFS98, BioXCell) or control rat IgG2a (BioXCell) both in 100 μl. Injection frequencies are shown in **Fig. S5**.

### Immunohistochemistry analysis

Tumors were fixed in 10% normal buffered formalin (VWR, Cat.No.: 16004-128) for 24 hrs, and further processed by the Northwestern University Mouse Histology & Phenotyping Laboratory (MHPL) (paraffin embedding, sectioning at 4 μm, slide preparation, and staining). Tissue sections of each specimen were stained using hematoxylin and Eosin (H&E) staining. The expression of F4/80, CD4, CD8, pSTAT1, CD31, FOXP3, Ly6G, CD11b, and NKp46 was determined in tumor and/or spleen slides. Paraffin sections were dewaxed and blocked with 5% BSA. The sections were then incubated with the relevant antibodies at 4°C overnight. The sections were washed with PBS.

For immunohistochemical analysis, sections were incubated with donkey anti-rabbit or donkey anti-rat antibody. Slides were examined and representative fields were photographed at 5x or 20x magnification using a Leica DM4000 B microscope. The immune-positive cells and staining intensities were quantified in a circle that was placed in each staining at the same position in a tumor of the 5x images. For each tumor and condition 8 fields were counted.

### IHC quantification

Stained slides on standard size (26 mm x 76 mm) were scanned using NanoZoomer 2.0-HT (020234 by Olympus America Inc.) with scanning mode of x20 (0.46 μm/pixel) either automatic or semi-automatic set up under SOP (MICR 1.1.03) at the department of Pathology and RHLCCC Core Facility of Northwestern University. The digital images were then stored as high definition and were analyzed with NDP (NanoZoomer Digital Pathology, Hamamatsu Inc.) View software (https://www.hamamatsu.com/us/en/product/type/U12388-01/index.html) for quantification analysis. For identification of individual nuclei in the tissue, we trained a Cell Classifier in the Visiopharm Image analysis Suite (VIS, Horsholm, Denmark) by manually annotating exemplar nuclei that were positive and negative for DAB (DAB+ and DAB-, respectively). Once the Cell Classifier was trained, we applied the trained Cell Classifier to all images to generate counts of DAB+ and DAB-nuclei and calculate a percent positive index. Three sets of tumors and two sets of spleens were analyzed for Ly6G, CD11b and TUNEL staining; in set 1 the entire tumor of multiple slides from 4T1 wt and CD95 k.o. in BALB/c mice were scanned, average percent positive cells for Ly6G and CD11 were determined. In set 2, either the entire tumor or 3 selected independent fields from pure tumor in multiple slides were quantified as average percent positive cells for TUNEL stain from 4T1 wt and CD95 k.o. in BALB/c mice treated with IgG or anti-CD8 Ab. Set 3 included the quantification of Ly6G staining in the entire tumor of multiple slides from 4T1 wt and CD95 k.o. tumors in NSG mice. Set 4 was the quantification of CD8, CD4, or NKp46 positive cells in spleens of depleted mice.

### RNA-seq library construction from cell lines grown *in vitro*

Total RNA from cells (U-clones) was extracted using TRIzol (Invitrogen 15596018). mRNAs were enriched using Oligo d(T)25 Magnetic Beads (NEB S1419S) following manufacturer’s protocol. mRNAs were then fragmented at 94 °C in 10mM MgCl2 buffer. Fragmented RNAs were end-repaired by T4 PNK (NEB M0201S) and poly(A)-tailed by E. coli Poly(A) Polymerase (NEB M0276S). The poly(A) tailed RNA fragments were reverse transcribed to cDNA using custom-designed oligo(dT) and locked nucleic acid (LNA) based on SMART-seq2 method [62]. The cDNAs were then PCR amplified using KAPA HiFi HotStart master mix (KK2601). 200-400 bp fragments of the library were cut from gels for Illumina Hiseq sequencing.

### RNA-seq library construction from tumors grown *in vivo*

For the RNA sequencing wt and CD95 k.o. 4T1 tumors were isolated from NGS and Balb/c mice and tumor tissues were store in RNA later solution until further use. For RNA isolation tumor tissue was lysed using Qiazol and total RNA was isolated using the miRNeasy Mini Kit (Qiagen, Cat.No# 74004) following the manufacturer’s instructions. An on-column digestion step using the RNAse-free DNAse Set (Qiagen, Cat.No# 79254) was included for all RNA-Seq samples. RNA libraries were generated and sequenced (Genomics Core at Northwestern University). The quality of reads, in FASTQ format, was evaluated using FastQC. Reads were trimmed to remove Illumina adapters from the 3’ ends using cutadapt [63]. Trimmed reads were aligned to the mouse genome (m38/mm10) using STAR [64]. Read counts for each gene were calculated using htseq-count [65] in conjunction with a gene annotation file for m38 obtained from Ensembl (http://useast.ensembl.org/index.html). Normalization and differential expression were calculated using DESeq2 that employs the Wald test [66]. The cutoff for determining significantly differentially expressed genes was an FDR-adjusted p-value less than 0.05 using the Benjamini-Hochberg method.

### RNA-seq and gene ontology data analysis

We used the first end of the sequencing reads for the analyses. The raw reads were filtered to remove low-quality reads using FASTQ Quality Filter from FASTX-Toolkit v0.0.14 (RRID:SCR_005534). High-quality reads were then mapped to the mouse reference genome (mm10) using STAR v2.6.0 (RRID:SCR_015899) with the following options to generate BAM files, count files, and read coverage tracks: --outSAMtype BAM SortedByCoordinate --quantMode TranscriptomeSAM GeneCounts –outWigType bedGraph –outWigNorm RPM. The transcriptomic BAM file was used to quantify gene expression using RSEM v1.3.0 (RRID:SCR_013027). Differential gene expression analysis was carried out on the gene counts using DESeq2 (RRID:SCR_015687). Gene expression (log2(TPM + 1)) was normalized to the mean of the corresponding control samples and significantly differentially expressed genes (DESeq2 adjusted p-value < 0.05) were clustered into 6 groups using k-means clustering. Gene ontology analysis was performed for each cluster using the TopGO package (RRID:SCR_014798). Heatmaps were created with the pheatmap package v1.0.12 (RRID:SCR_016418). All analyses were carried out using R v3.6.3 (RRID:SCR_001905). The GO enrichment analyses shown in Fig. 3B was performed using the GOrilla gene ontology analysis tool at http://cblgorilla.cs.technion.ac.il using default settings and a p-value cut-off of 10^-6^.

## Statistical methods

All experiments were performed in triplicate. The results were expressed as mean ± SD and analyzed by the Student’s two-tailed t test or by ANOVA (Prism8). Statistical significance was defined as p < 0.05.

## Data availability

Sequencing data have been deposited in the National Cancer for Biotechnology Information Gene Expression Omnibus with accession number GSE154676 (for the RNAseq analysis of the cell lines) and GSE154682 (for the RNAseq data of the tumors grown in mice).

## Acknowledgement

U-clones: We would like to thank Shanshan Zhang for performing slide scanning and signal quantification and Michelle von Locquenghien for help with the generation of the 4T1 CD95 k.o. clones. F-clones: We would like to thank the IHC facility (Dr. Alain Fautrel and Roselyne Viel, Rennes, France) for performing 4T1 tissue staining, slide scanning and signal quantification. The mouse study in NSG mice benefited from the help of Remy Castellano and Emmanuelle Josselin (Marseille, France). This work was funded by grant R35CA197450 to M.E.P, by INCa (PLBIO18-059), Ligue Contre le Cancer and Fondation de France (Price Jean Valade) for P.L. and by Lynn Sage Cancer Research Foundation and Lynn Sage Scholar fund for Z.J.

## Author contributions

A.S.Q., J.P.G., C.G., E.C-J, A.B., V.L., T. M-R., P.L., Z.J. and M.E.P. planned the experiments. A.S.Q., J.P.G., C.L., H.B., E.S., A.C., J.V., M.J.S., M.M., A.E.M, and B.B. performed experiments or analyzed data. P.L. and M.E.P. directed the study and wrote the manuscript.

