## Supplemental figures for "Tumor expressed CD95 causes suppression of anti-tumor activity of NK cells in a model of triple negative breast cancer"

**Appendix files - Table of contents**

**Appendix Figures:**

**Figure S1.** Generation and *in vitro* characterization of two sets of CD95 k.o. 4T1 cells................ p2

**Figure S2.** Unresponsiveness of 4T1 cells to CD95 stimulation.................................................... p4

**Figure S3.** Deletion of CD95 does not consistently change cancer stemness in 4T1 cells............. p5

**Figure S4.** TAMs infiltrate wt 4T1 tumors in NSG mice better than CD95 k.o. tumors................ p6

**Figure S5.** Treatment schemes of immune cell depletion experiments........................................... p7

**Figure S6.** Lack of infiltration of $CD8^+$ T cells into 4T1 tumors with deleted CD95 in Balb/c mice after depletion of CD8.......................................................................................... p8

**Figure S7.** Depletion of CD4 T cells or of macrophages does not reverse reduced growth of CD95 k.o. tumors in Balb/c mice............................................................................................ p9

**Table S1:** Source data for Figure 3A

**Table S2:** Source data for Figure 3B

**Table S3:** Subset of genes in Table S1 that are either CSF ligands or receptors

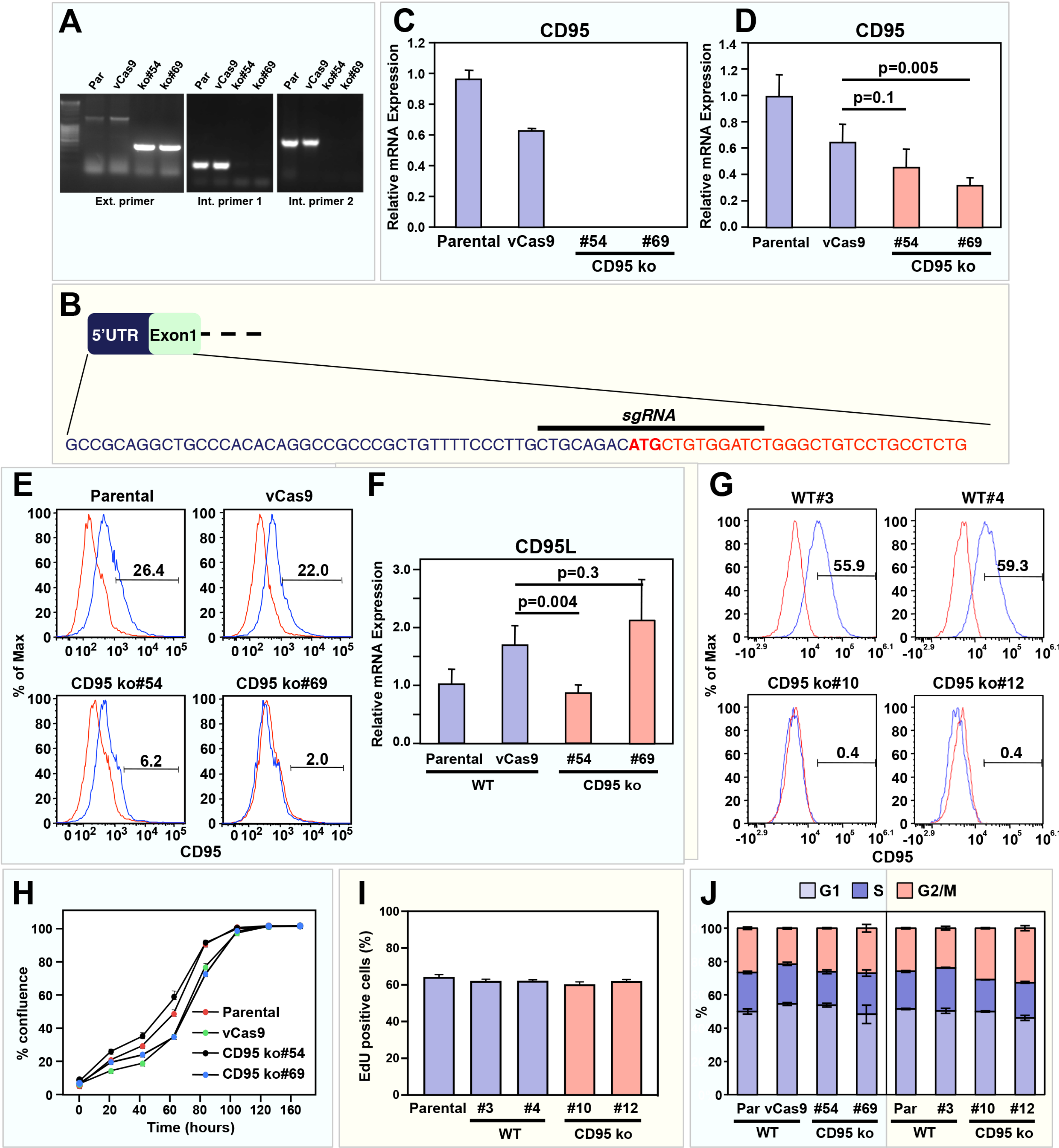
A
Par
vCas9
ko#54
ko#69
Ext. primer
Int. primer 1
Int. primer 2
C
CD95
Relative mRNA Expression
Parental
vCas9
#54
#69
CD95 ko
D
CD95
p=0.005
p=0.1
B
5'UTR
Exon1
sgRNA
GCCGCAGGCTGCCCACACAGGCCGCCCGCTGTTTTCCCTTGCTGCAGACATGCTGTGGATCTGGGCTGTCCTGCCTCTG
E
Parental
vCas9
26.4
22.0
CD95 ko#54
CD95 ko#69
6.2
2.0
% of Max
CD95
F
CD95L
p=0.3
p=0.004
WT
CD95 ko
G
WT#3
WT#4
55.9
59.3
CD95 ko#10
CD95 ko#12
0.4
0.4
H
% confluence
Parental
vCas9
CD95 ko#54
CD95 ko#69
Time (hours)
I
EdU positive cells (%)
#3
#4
#10
#12
J
G1
S
G2/M
%
Par
vCas9

**Appendix Figure S1. Generation and *in vitro* characterization of two sets of CD95 k.o. 4T1 cells**.

(**A**) Characterization of Parental, vCas9 clone and two exon9 specific deleted CD95 k.o. clones of 4T1 cells. Genotyping of homozygous deletions exon 9 of CD95 in two isolated U-clones #54 and #69 documented by with one external and two different internal primer pairs.

(**B**) Schematic representation of the CD95-targeting sgRNA for the generation of F-clones.

(**C, D, F**) Real time PCR analysis of parental, a vCas9 clone and two CD95 k.o. clones of 4T1 with a mouse CD95 exon 9 specific primer (C), a mouse CD95 exon 1-2 specific primer (D), or CD95L specific primer (F).

(**E**) CD95 surface staining of parental, a vCas9 clone and the two CD95 k.o. U-clones of 4T1 cells.

(**G**) CD95 surface expression in two wt and two CD95 k.o. 4T1 F-clones analyzed by flow cytometry (**H**) Cell growth over time of the same cells.

Data are representative of two independent experiments. Each data point represents mean ±SE of three replicates.

(**I**) Indicated 4T1 F-clones were incubated with the thymidine analog EdU, then washed and fixed. By 'click' chemistry, iFluor-488 was covalently conjugated to the EdU and the percentage of cells in S-phase (EdU positive cells) was assessed by flow cytometry. Data represent mean ±SD of three independently performed experiments.

(**J**) Cell cycle analysis of exponentially growing U and F parental, wt and CD95 k.o. clones.

p-value *<0.05, **<0.001; ***<0.0001; ns, not significant.

Experiments involving U-clones are in a light blue box and experiments involving F-clones are in a light yellow box, respectively.

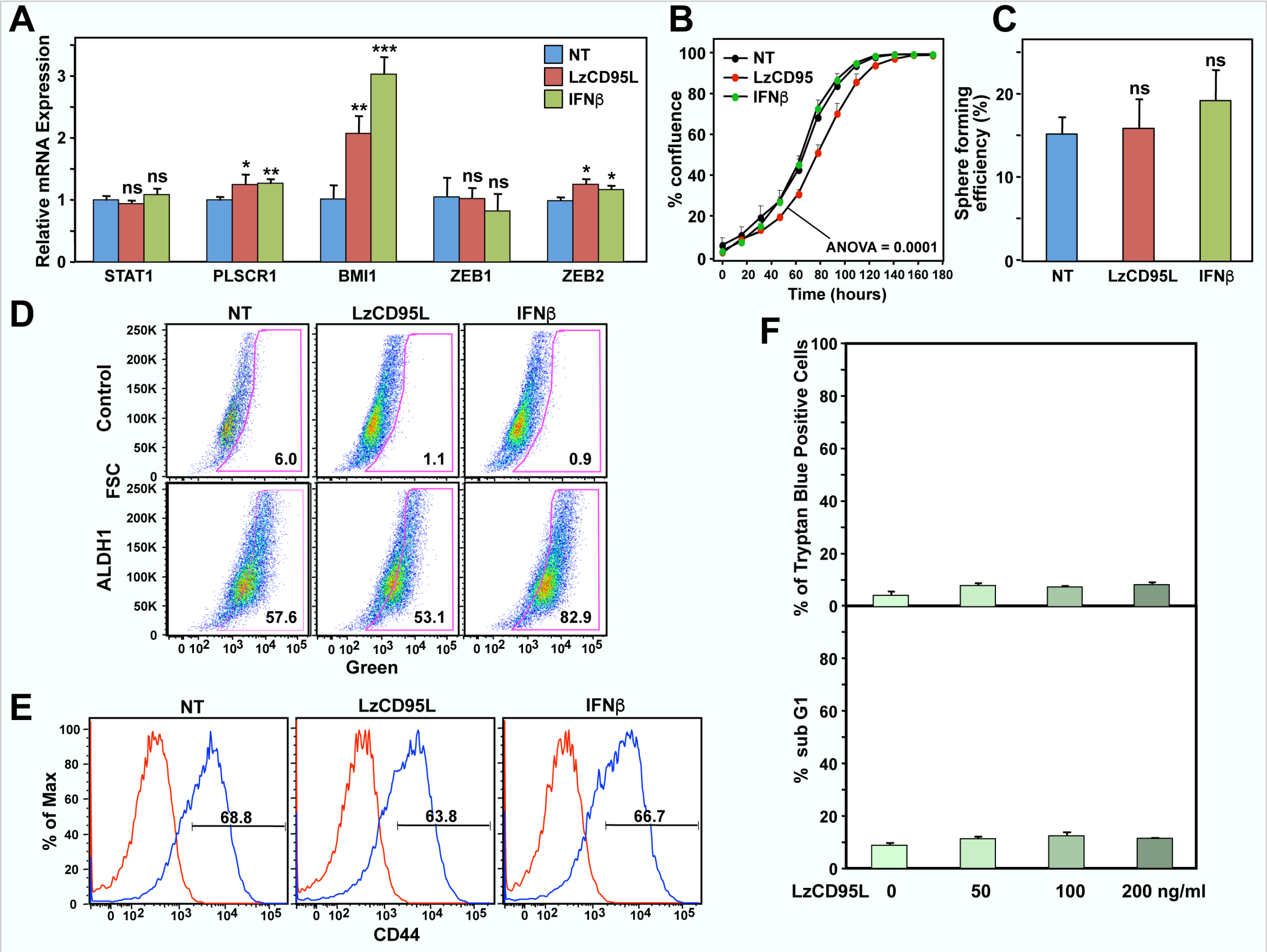


**Appendix Figure S2: Unresponsiveness of 4T1 cells to CD95 stimulation.**

(**A**) Real-time PCR analysis of STAT1, the STAT1 target gene PLSCR1 and stem cell marker genes BMI1, ZEB1, and ZEB2 in 4T1 cells after a 4-day treatment with CD95L or IFNβ.

(**B**) Growth of the 4T1 cells untreated or treated with either LzCD95L or IFNβ. p-value was calculated using ANOVA.

(**C**) Single cell sphere formation assay of 4T1 cells untreated or treated with either LzCD95L or IFNβ for 6 days.

(**D**) ALDH1 activity of 4T1 cells untreated or treated with either LzCD95L or IFNβ for 6 days.

(**E**) CD44 surface staining of 4T1 cells untreated or treated with either LzCD95L or IFNβ for 6 days.

(**F**) Percent of nuclear PI staining (left panel) or percent of Trypan blue positivity (right panel) of 4T1 cells 24 hrs after adding different amounts of LzCD95L.

p-value *<0.05, **<0.001; ***<0.0001; ns, not significant. Experiments were performed with the U-clones.

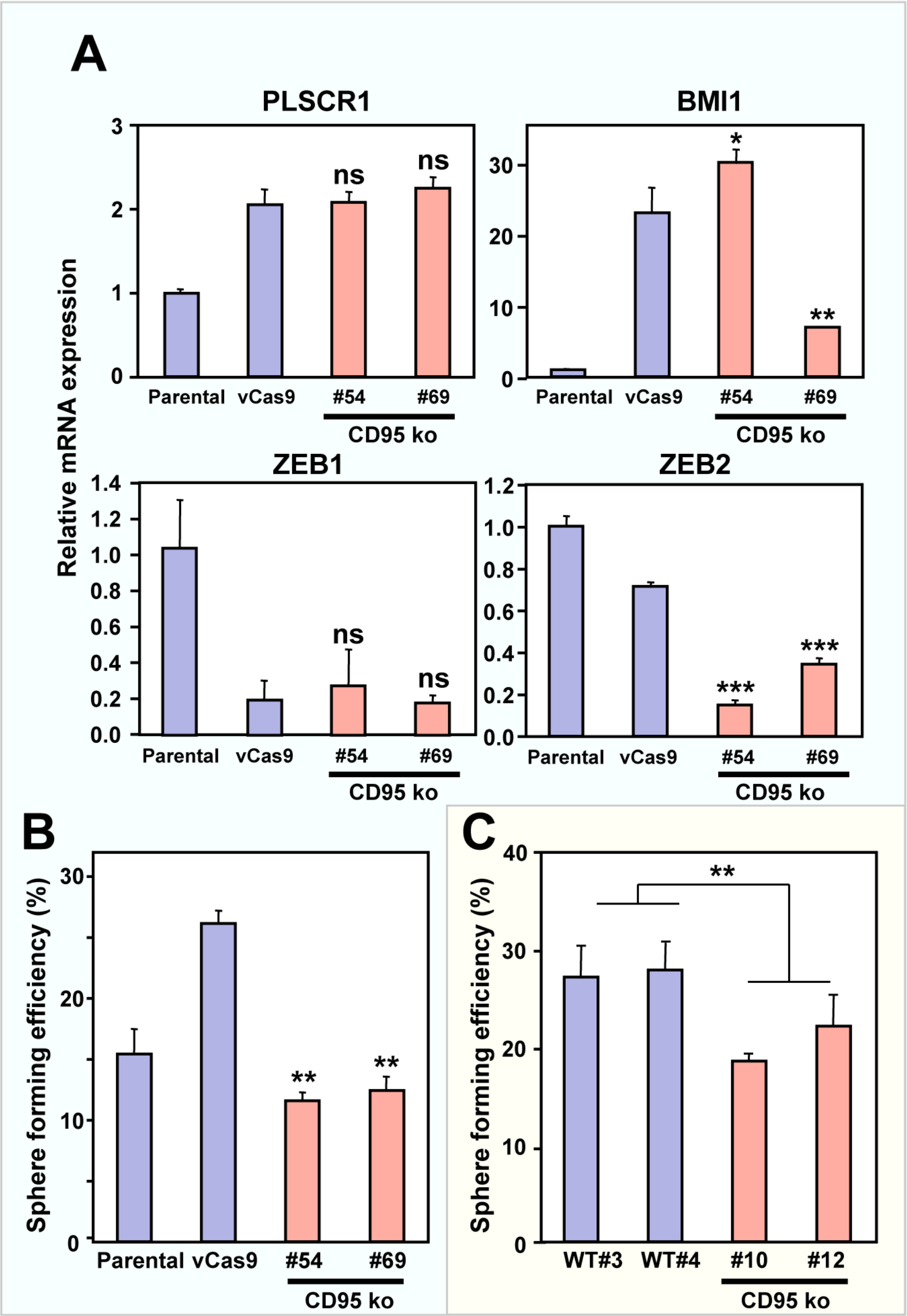


**Appendix Figure S3. Deletion of CD95 does not consistently change cancer stemness in 4T1 cells.**

(**A**) Real-time PCR analysis of PLSCR1 and stem cell marker genes BMI1, ZEB1 and ZEB2 of parental, vCas9 clone and two CD95 k.o. clones.

(**B**) Single cell sphere formation assay of parental, vCas9 clone and two CD95 k.o. U-clones. p-value *<0.05, **<0.001; ***<0.0001; ns, not significant.

(**C**) Single cell sphere formation assay of two wild type and two CD95 k.o. F-clones. Mann-Whitney test p=0.0065.

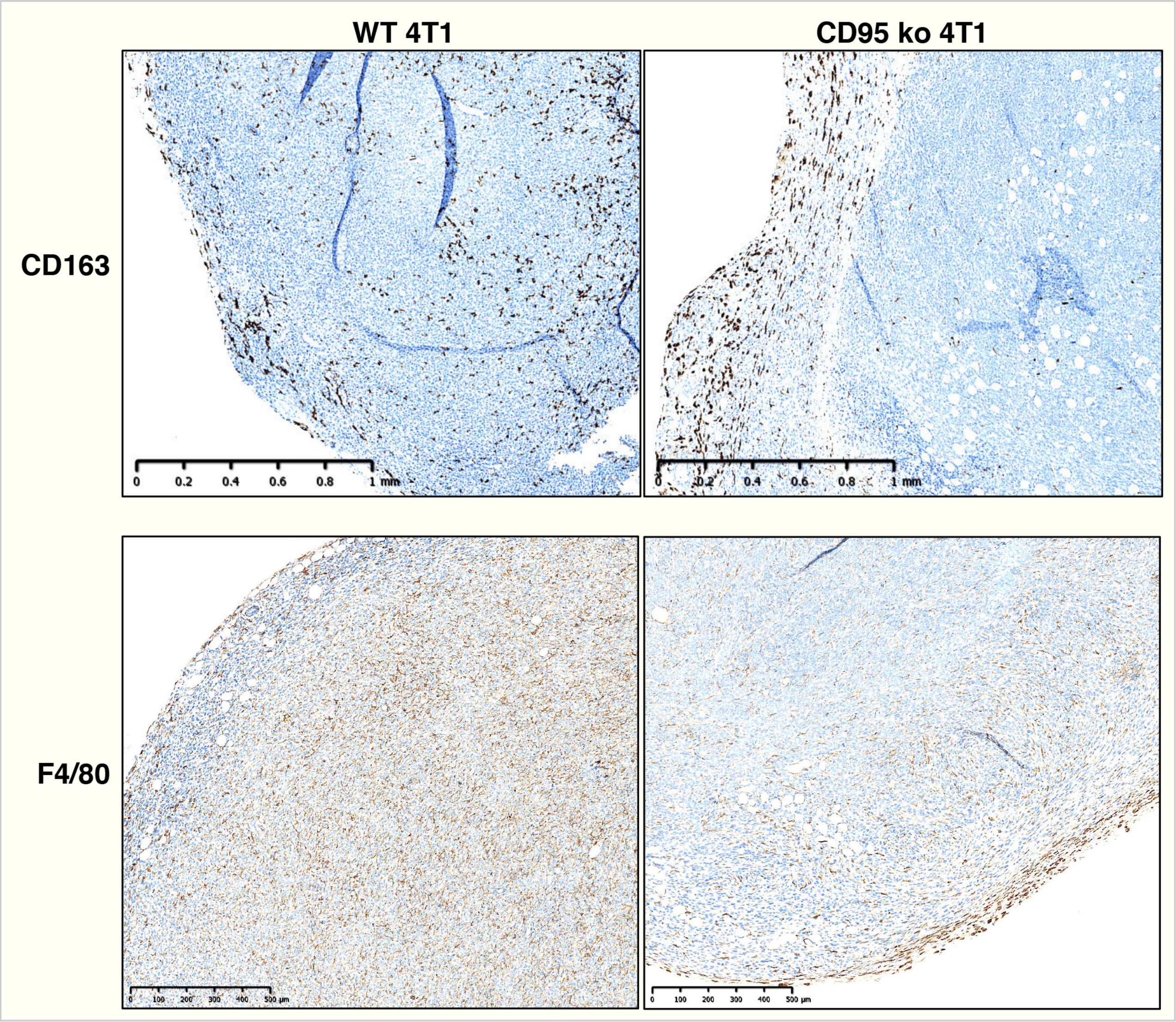


**Appendix Figure S4. TAMs infiltrate wt 4T1 tumors in NSG mice better than CD95 k.o. tumors.** Representative immunohistochemical staining of TAMs in wt (clone #3) and 4T1 k.o. (clone #12) F-clones.

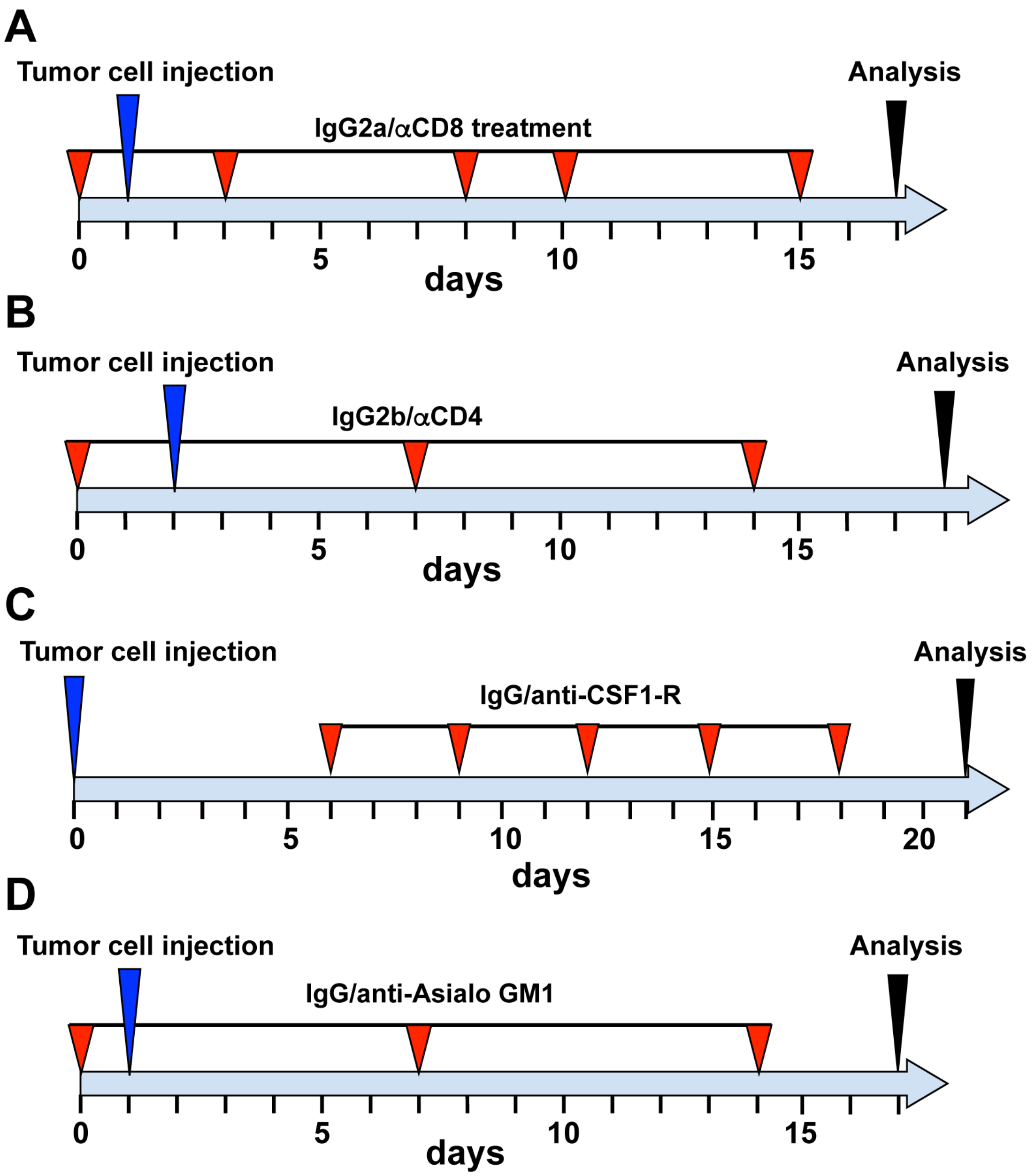


**Appendix Figure S5. Treatment schemes of immune cell depletion experiments.**
Treatment scheme of Balb/c with implanted wt or CD95 k.o. cells and treated with control Ab or andi-CD8 (A), anti-CD4 (B), anti-CSF1-R (C), or anti-Asialo GM1 (D) antibodies.

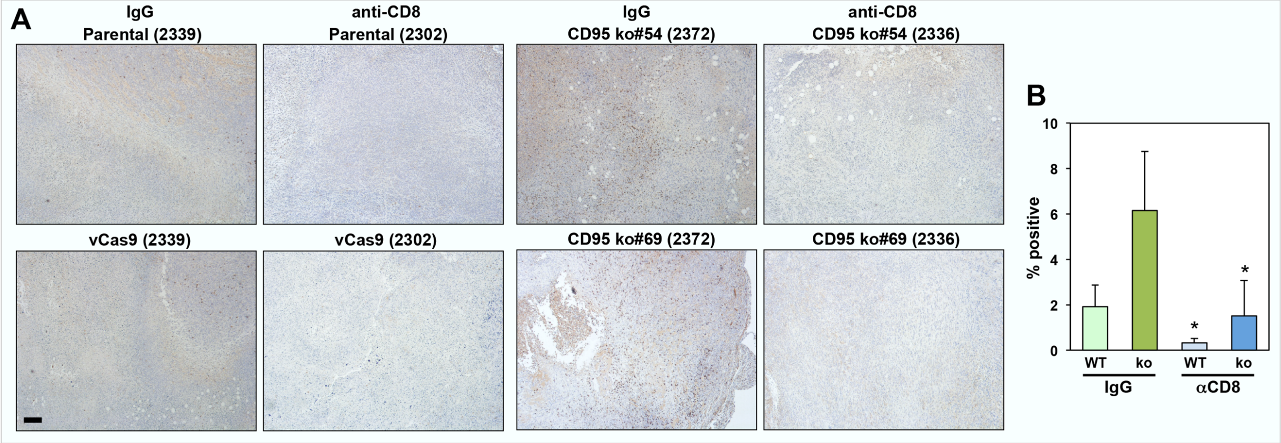


**Appendix Figure S6. Efficiency of the CD8 depletion in the tumor of 4T1-grafted Balb/c mice.**
(**A**) IHC of wt or CD95 k.o. tumors grown in Balb/c mice after depletion of $CD8^+$ T cells staining with anti-CD8 mAb. Mouse ear tag numbers are shown in brackets. Experiments were performed with the U-clones. Scale bar = 50 µm.
(**B**) Quantification of CD8 T cells in tumors of mice treated with either IgG or anti-CD8 mAb. p-value *<0.05.

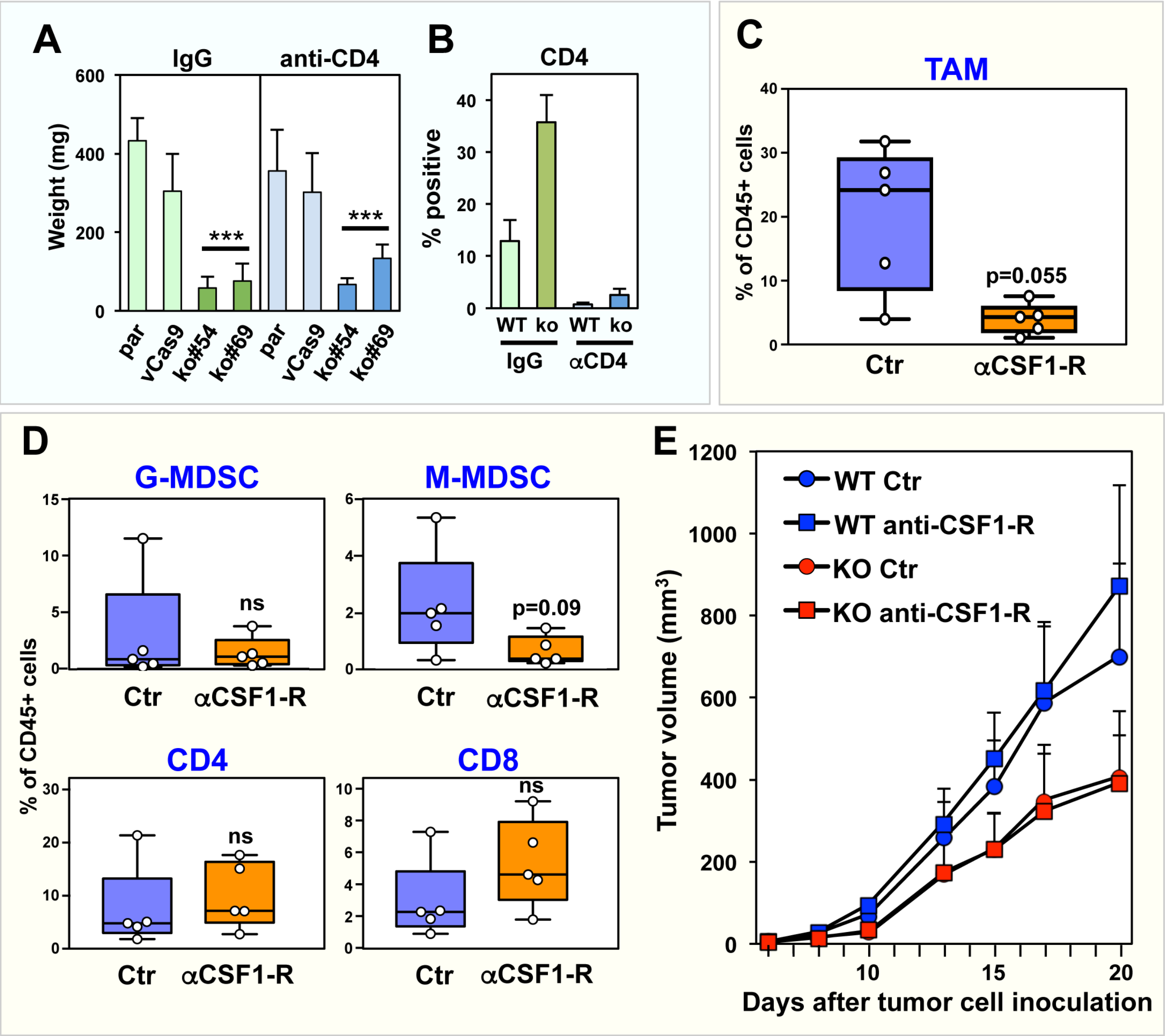


**Appendix Figure S7. Depletion of CD4 T cells or of myeloid cells does not reverse reduced growth of CD95 k.o. tumors in Balb/c mice.**

(**A**) Tumor weight (left) and tumor volume (right) in anti-CD4-treated mice (n=5) 16 days after tumor injection.

(**B**) Quantification of CD4 T cells in the spleen of mice treated with either IgG or anti-CD4 mAb.

(**C**) Multiparameter flow cytometry quantification of tumor-infiltrating macrophages ($CD45^{+}CD11b^{+}F4/80^{+}$) in mice treated with either control IgG or anti-CSF1-R mAb, 21 days after 4T1 cell injection. Mann-Whitney test was used to calculate p-value.

(**D**) Multiparameter flow cytometry quantification of tumor-infiltrating G-MDSC ($CD45^{+}CD11b^{+}Ly6C^{Low}Ly6G^{+}$), M-MDSC ($CD45^{+}CD11b^{+}Ly6C^{+}$Ly6G-), and $CD3^{+}CD4^{+}$, $CD3^{+}CD8^{+}$ T-cells in mice treated with either control IgG or anti-CSF1-R mAb, 21 days after tumor injection.

(**E**) Tumor growth of wt (clone #3) or CD95 k.o. (clone #12) clones in mice treated with either control or anti-CSF1-R antibody.
